# Photoproximity Profiling of Protein-Protein Interactions in Cells

**DOI:** 10.1101/833384

**Authors:** David C. McCutcheon, Gihoon Lee, Anthony Carlos, Jeffrey E. Montgomery, Raymond E. Moellering

## Abstract

We report a novel photoproximity protein interaction (PhotoPPI) profiling method to map protein-protein interactions *in vitro* and in live cells. This approach utilizes a bioorthogonal, multifunctional chemical probe that can be targeted to a genetically encoded protein of interest (POI) through a modular SNAP-Tag/benzylguanine covalent interaction. A first generation photoproximity probe, PP1, responds to 365 nm light to simultaneously cleave a central nitroveratryl linker and a peripheral diazirine group, resulting in diffusion of a highly reactive carbene nucleophile away from the POI. We demonstrate facile probe loading, and subsequent interaction- and light-dependent proximal labeling of a model protein-protein interaction (PPI) *in vitro*. Integration of the PhotoPPI workflow with quantitative LC-MS/MS enabled unbiased interaction mapping for the redox regulated sensor protein, KEAP1, for the first time in live cells. We validated known and novel interactions between KEAP1 and the proteins PGAM5 and HK2, among others, under basal cellular conditions. By contrast, comparison of PhotoPPI profiles in cells experiencing metabolic or redox stress confirmed that KEAP1 sheds many basal interactions and becomes associated with known lysosomal trafficking and proteolytic proteins like SQSTM1, CTSD and LGMN. Together, these data establish PhotoPPI as a method capable of tracking the dynamic sub-cellular and protein interaction “social network” of a redox-sensitive protein in cells with high temporal resolution.

**SYNOPSIS TOC:** 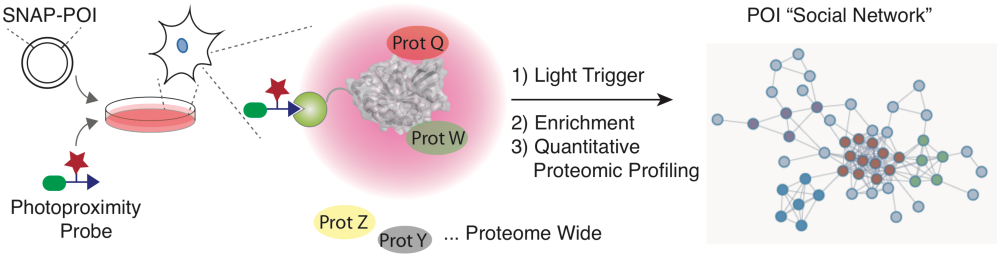

## INTRODUCTION

A grand challenge in biology is to build high-fidelity and dynamic maps of molecular interaction networks inside of living cells. The ability to elucidate the so-called “social network” for a biomolecule of interest under a wide range of biological contexts would provide the basis for understanding how cellular machinery is organized in space and time^1^. This information would represent a quantum leap forward in our understanding of cellular signaling events, as well as the potential to perturb these events for therapeutic purposes. To draft high-quality interactome maps we must develop methods that can detect transient binding interactions, ideally while preserving the spatial, mechanical, and chemical environment found in living systems (i.e. cells or tissues). Affinity purification-mass spectrometry (AP-MS) pull-down approaches have historically been used to enrich a tagged protein-of-interest (POI), with the intention that binding partners will co-elute during iterative rounds of enrichment and washing, followed by protein detection by mass spectrometry^2^. While powerful, the steps involved in cellular lysis and dilution through iterative washes biases against weaker binding interactions that dominate many protein-protein interactions, and also has the propensity to introduce non-specific interactions between partners that may have been compartmentalized in the cell. There are many classes of interactions and proteins that are not appropriate for this approach, including membrane proteins, chromatin associated complexes, and redox-sensitive complexes, among others. Therefore, while many of the protein interactome maps drafted to date have been developed with AP-MS methods, new methods to augment or replace these maps are needed.

To circumvent the need to lyse cells and perform *in vitro* enrichments, several new technologies have been developed that enable chemical or enzymatic tagging of proximal protein binding partners within live cells, which are then retrieved after cellular lysis and identified by LC-MS/MS. The BioID method fuses an engineered biotin ligase *BirA* to a target POI, which then converts intracellular ATP and exogenous biotin to an amine-reactive biotinoyl-5’-AMP that can diffuse from the POI and covalently label proximal proteins^3-4, 5^. Proximal enzymatic labeling with similarly reactive thioesters, for example with the NEDDylator^6^, PUP-IT^7^, and EXCEL^8^ systems, have been shown to label proximal proteins with small protein or peptide tags, enabling subsequent profiling. Genetic fusions of horseradish peroxidase^9^ and engineered ascorbic acid peroxidase (i.e. APEX) enzymes can convert exogenous chemical probes into reactive phenoxyl radicals^10-11^ to label the sub-cellular proteome in organelles^11-12^, as well as identify protein complex members in diverse conditions^13^. Even with the success of these approaches, there are significant limitations for specific applications imposed by the requirement for co-factors, an exogenous biotin-phenol probe, and high levels of H_2_O_2_ to initiate labeling, which could bias efficient labeling in specific cellular and chemical environments. Additionally, the requirement for peroxide limits application of this approach with proteins or pathways that involve redox regulation, which likely includes a large percentage of the proteome^14-16^. Therefore, new methods that can label proximal proteins in live cells with high spatial and temporal control, ideally without significant perturbation to the cellular environment, would provide a significant advantage to mapping PPIs. Herein we present a novel photoproximity protein interaction (PhotoPPI) profiling platform that uses light to trigger proximal protein labeling in live cells. We demonstrate protein-interaction-dependent covalent labeling of binding partners with this approach *in vitro*, as well as proteome-wide identification of steady state binding partners for the highly redox sensitive protein KEAP1, which validated a network of known and novel protein interactors perturbed metabolism or redox status.

**Scheme 1.**
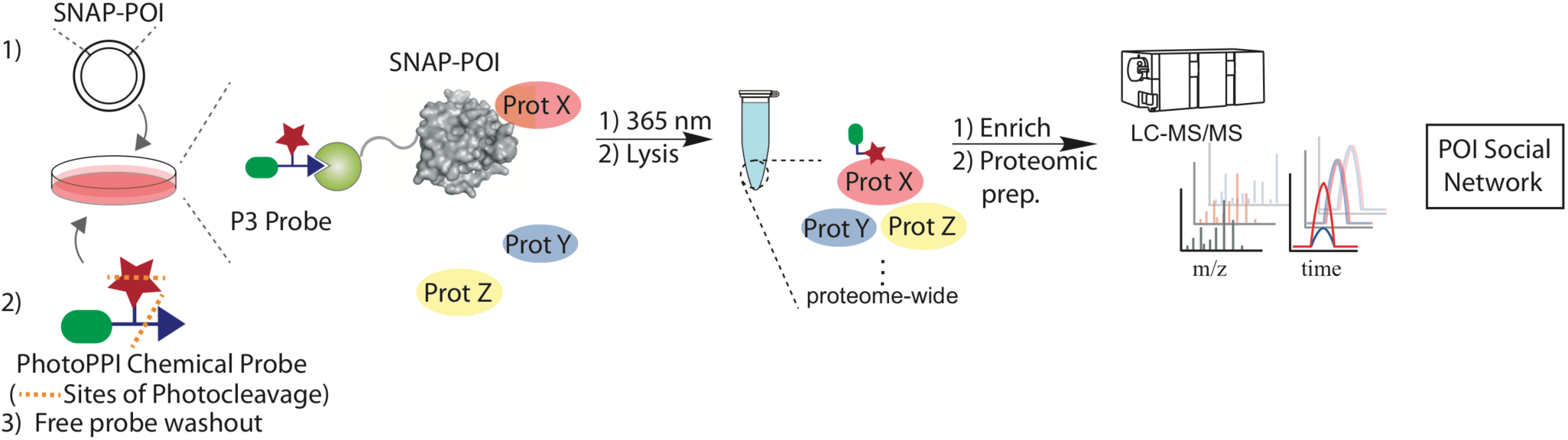
PhotoPPI profiling workflow. Photoproximity probe treatment and subsequent washout of SNAP-POI-expressing cells results in *in situ* tagging of POI proteins. Subsequent light-triggered, dual photocleavage events (depicted with orange dashes on probe structure) releases a reactive carbene to covalently label proximal proteins. Enrichment of labeled proteins and quantitative analysis by LC-MS/MS can identify the proximal “social network” for a protein of interest.

## RESULTS AND DISCUSSION

### Design of the PhotoPPI platform

We designed a first generation PhotoPPI chemical proteomic method to facilitate several activities, including: 1) specific probe labeling of POIs in live cells; 2) spatial and temporal control of probe photoactivation and proximal labeling; 3) subsequent enrichment and identification of labeled proteins after cell lysis (Scheme 1). To realize these requirements, we first enabled targeting of our modular photoproximity chemical probe, PP1, to a POI through a benzylguanine (BnG) recognition element, which has been shown to specifically label a “SNAP-Tag” protein (an engineered *O*^*6*^*-*methylguanine DNA methyltransferase, MGMT) that can be theoretically expressed as a genetically-encoded fusion on any POI (Scheme 1; Fig. 1A)^17-18^. The BnG targeting element is connected to dual photoreactive elements, a central substituted nitroveratryl carbamate, and a tethered diazirine, which upon irradiation with 365 nm light will simultaneously trigger cleavage and diffusion of the probe away from the SNAP-Tag-POI, as well as unmasking of a highly reactive carbene, respectively (Fig. 1A). Finally, the diazirine portion of PP1 is connected to a retrieval tag - in this case biotin - for recognition and enrichment of proteins that were covalently labeled in live cells (Fig. 1A). We first validated the labeling and photocleavage capacity of the central nitrobenzyl carbamate linker using a cleavable BnG-FITC model probe (PF-BnG), which fluorescently labeled recombinant SNAP-Tag protein (Fig. 1B). In line with other reports using photocleavable linkers for selective delivery of bioactive molecules on or in cells^19-20^, we found that irradiation of the labeled protein at 365 nm resulted in complete loss of fluorescence within minutes, validating the capacity to label and cleave BnG-nitrobenzyl probes. In parallel, we tested the photo-cleavage kinetics of the nitroveratryl and diazirine portions of PP1 in bulk solution by LC-MS, which confirmed half-lives of just under a minute (Suppl. Fig. 2). To directly test the localization and light-dependent cleavage of a probe from SNAP-Tag in cells, we treated cells expressing a nuclear-localized SNAP construct (SNAP-FLAG-NLS) with PF-BnG, which resulted in localized fluorescent signal in the nucleus (Fig. 1C; Suppl. Fig. 3A-C). Irradiation at 365 nm led to rapid loss of fluorescence after approximately one minute, in agreement with the *in vitro* kinetics studies (Suppl. Fig. 3c). To determine whether the multifunctional photoprobe PP1 probe could label SNAP-Tag protein in cells, we treated HEK293T cells stably expressing FLAG-tagged SNAP-Tag protein (SNAP-FLAG) with increasing doses of PP1 for two hours, which resulted in robust covalent labeling at low micromolar concentrations (EC_50_ = 4.5 μM; Fig. 1D). Importantly, cell viability was not affected at saturating doses of PP1 or UV irradiation (Fig. 1E-F), suggesting that cells expressing a SNAP-Tag-POI fusion could be pulse-labeled with PP1 with minimal perturbation to cell physiology.

**Figure 1.**
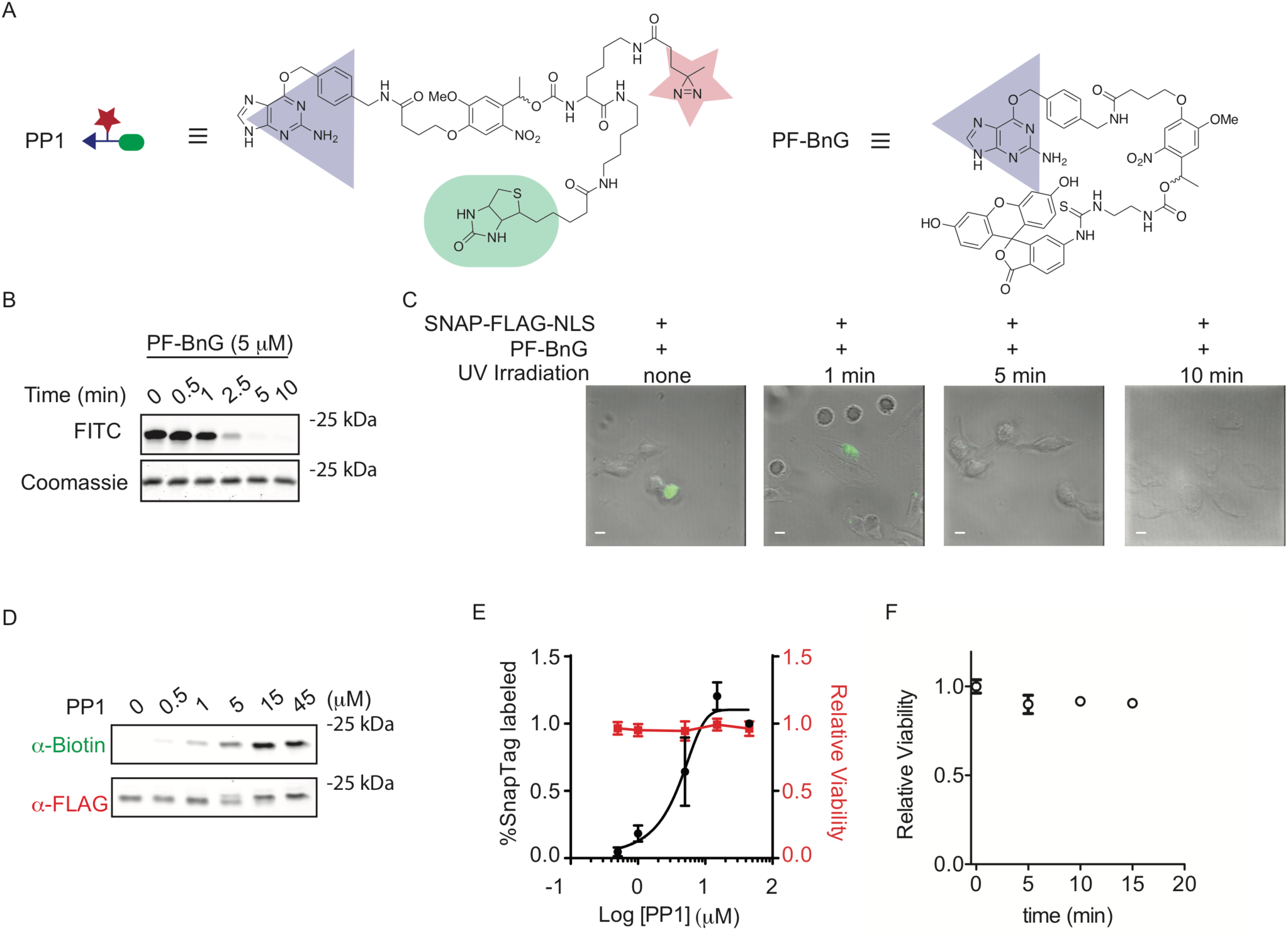
Photoproximity probe labeling and photocleavage in cells. A) Chemical structures and design elements of the first generation photoproximity probe 1, PP1, and the fluorescent model compound, PF-BnG. B) Fluorescence gel-based visualization of recombinant SNAP protein labeled with the model probe PF-BnG, exposed to UV light for the indicated time *in vitro*.C) Confocal fluorescence images of HeLa cells transfected with SNAP-FLAG-NLS treated with PF-BnG and exposed to UV light for the indicated time. Fluorescence and DIC overlays are shown for a representative field. Full views are in Suppl. Fig. 3D) Western blot detection of total and biotin-labeled SNAP-FLAG protein expressed in HEK293T cells and treated with indicated doses of PP1 for two hours. Cells and lysates were not irradiated prior to gel and Western blot. E) Quantification of SNAP-FLAG protein labeling and HEK293T cell viability (determined by relative ATP content) at indicated PP1 doses. F) HeLa cell viability after exposure to 365 nm light for the indicated timepoints as measured by cell titer glo assay. Gels and Western blots shown are representative from two or more biological replicates. Points and error bars represent the mean and standard error of the mean from two or more biological replicates. Scale bars in (C) = 8.3 μm.

### *In vitro* protein-protein interaction profiling

To test if proximal protein interactors could be labeled in response to the dual photoactivation of PP1 *in vitro*, we examined the interaction between SNAP-FLAG protein and an *α*-FLAG monoclonal antibody (mAb; Fig. 2A) in cell lysate. In this assay, we would expect proximal biotinylation of both heavy and light chains of the antibody, as well as self-labeling of the SNAP-FLAG protein under conditions where the protein complex is formed and the PP1 probe is photoactivated (Fig. 2A). To test this hypothesis in the presence and absence of the protein-protein complex, we treated SNAP protein that contained or lacked the FLAG epitope with PP1 probe, followed by incubation with *α*-FLAG mAb, bead-based pulldown of the complexes, and subsequent photoactivation. Anti-mouse (recognizing both heavy and light chain *α*-FLAG) and streptavidin-IR western blot revealed robust biotinylation of both heavy and light chain fragments with SNAP-FLAG protein, but no protein-biotinylation in reactions using SNAP protein that lacked the FLAG sequence, validating the required protein-complex for proximal labeling by PP1 (Fig. 2B). We further tested the requirement for photoactivation of PP1, as well as a related probe PP2 (Suppl. Fig. 1), by forming the SNAP-FLAG/*α*-FLAG complexes, and then comparing protein biotinylation with or without irradiation. Complexes labeled with PP1 or PP2, but not exposed to 365 nm light exhibited biotinylation of a protein at ∼ 20 kDa, consistent with SNAP protein by molecular mass, due to the retention of the active-site labeled probe. No proximal labeling of the heavy chain protein, or a closely migrating light chain protein was observed without irradiation (Fig. 2C). By contrast, a discrete biotinylated band was formed on both heavy and light chains upon treatment with and irradiation of both PP1 and PP2 (Fig. 2C). The same pattern of light-dependent heavy chain labeling was observed if the complex was formed in isolation (Fig. 2C) or in whole proteome (Suppl. Fig. 4). Importantly, the bead pulldown and washing of PP1-labeled SNAP protein in this assay mimics the ability to remove unbound probe by washout in cell culture, thus only activating the photoprobe on the SNAP-POI for specific proximity labeling. Together, these data confirmed the light- and proximity-dependent labeling of protein complex partners with the PhotoPPI system.

**Figure 2.**
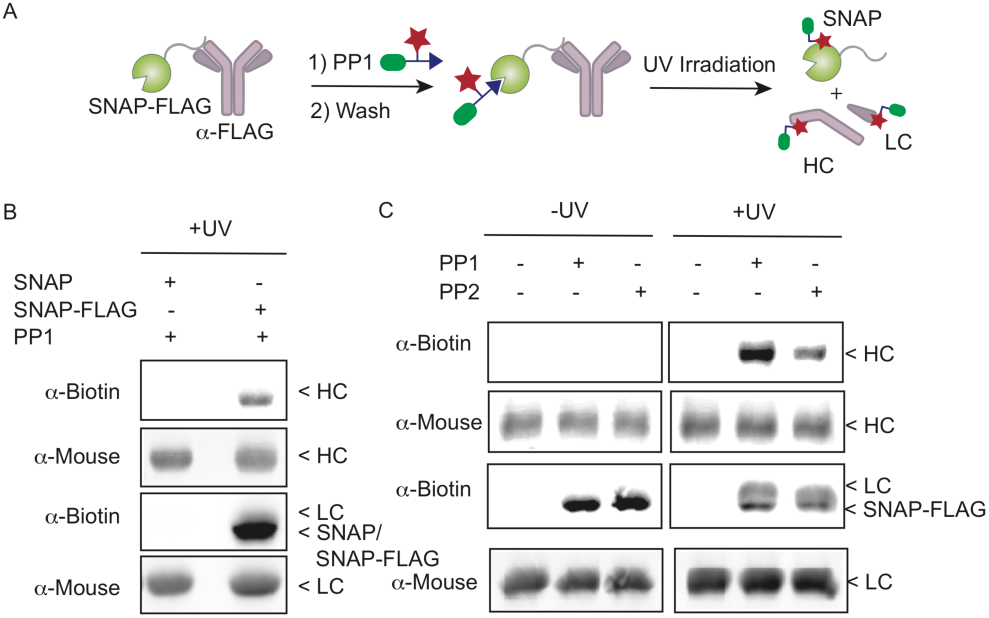
Proximity- and light-dependent protein complex labeling with photoproximity probes. A) Schematic depiction of the model protein-protein complex formed between SNAP-FLAG and *α*-FLAG antibody. Theoretical photoproximity labeling of individual proteins is shown. ‘Wash’ briefly refers to bead-based pulldown and removal of un-bound probe, which is analogous to the cell-based profiling workflow (*vide infra*)B) Anti-biotin (streptavidin-800) and anti-mouse Western blot analysis of PP1 labeled SNAP and SNAP-FLAG protein incubated with *α*-FLAG antibody prior to irradiation. C) Anti-biotin (streptavidin-800) and anti-mouse Western blot analyses of PP1 labeled SNAP-FLAG/ *α*-FLAG antibody complex with and without UV irradiation prior to analysis. Labels for individual proteins are included at appropriate molecular weights: LC, light chain; HC, heavy chain; “SNAP” label represents SNAP-Tag protein without the FLAG epitope.

### Photoproximity profiling in live cells

We next employed the PhotoPPI platform to determine whether proximal binding partners for a POI could be interrogated in live cells. In particular, we aimed to test this approach with a protein that would be challenging to study by existing proximity profiling methods. With these purposes in mind, we chose KEAP1, which acts as the critical sensor protein at the center of the antioxidant response signaling network.

KEAP1 harbors numerous reactive cysteines that collectively sense alterations in the redox, metabolic and xenobiotic environments of the cell; these interactions ultimately control the sequestration and turnover of the NFE2L2 (also known as NRF2) transcription factor and the downstream antioxidant gene expression program^21^. Given the central role of KEAP1 protein in sensing and controlling the cellular environment, a global assessment of its binding partners in specific cells and under discrete conditions could provide novel insights into core cellular wiring and physical interactions with other pathways.

We first cloned both C- and N-terminal SNAP-Tag fusions of full-length, human KEAP1, in each case separated by a short linker (Suppl. Fig. 5). Stable expression in HEK293T cells resulted in the appearance of the KEAP1-SNAP protein monomer at the expected mass of 90 kDa (Fig. 3A). We confirmed that the SNAP fusion was functional when expressed on KEAP1, as shown by dose-dependent labeling with PP1 in live cells (Fig. 3A). The low micromolar EC_50_ indicated that cellular studies with ∼15 μM PP1 would be suitable to label all fusion proteins prior to irradiation. Finally, we reasoned that identification of probe-labeled binding partners, and therefore proteins proximal to KEAP1 in cells, would be improved by quantitative proteomic differentiation of enriched vs. background proteins. We therefore SILAC-labeled cultures expressing both N-terminal (referred to as SNAP-KEAP1) and C-terminal (referred to as KEAP1-SNAP) fusion proteins with light and heavy arginine and lysine (Fig. 3B). Matched cultures of heavy and light cells, each expressing the same SNAP-KEAP1 or KEAP1-SNAP construct, were then treated with PP1 probe or vehicle for 2 hr. Following compound washout, cells were irradiated, lysed in denaturing RIPA buffer, and exposed to a streptavidin-bead pulldown of biotinylated proteins from each sample. Separate LC-MS/MS analysis of the combined bulk proteome, and combined heavy/light enriched proteome, was performed to selectively identify proteins that are enriched in the heavy, PP1 proximity labeled proteome, relative to any background effects in the bulk proteome. Together these profiles were integrated to identify the basal, proximal “social network” of KEAP1 in live cells (Fig. 3B).

**Figure 3.**
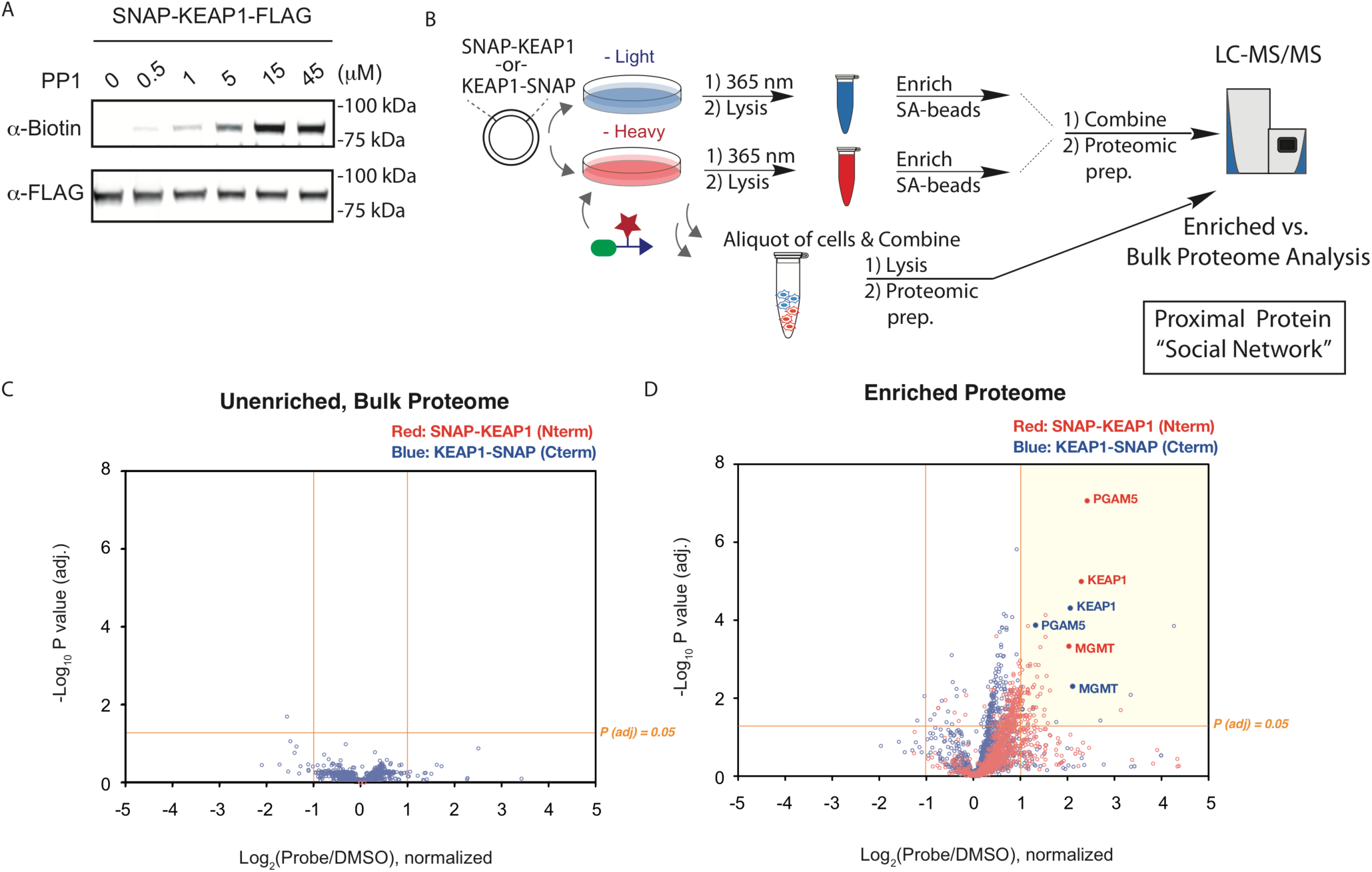
PP1 labels KEAP1 fusion proteins and labels proximal protein interactors in live cells. A) Anti-biotin (streptavidin-800) and anti-FLAG Western blot analysis of PP1 labeled KEAP1-SNAP protein from HEK293T cells treated with the indicated PP1 doses for 2 hr. Cells and lysates were not irradiated prior to gel analysis. B) Schematic of PhotoPPI profiling workflow using SILAC-labeled cells expressing SNAP-KEAP1 constructs. Both bulk and anti-biotin enriched proteome profiles are integrated to identify KEAP1 binders in cells. C) Volcano plot graph of bulk protein abundance SILAC ratios and *P*-values for both SNAP-KEAP1 (red) and KEAP1-SNAP (blue) expressing cells treated with PP1 probe (heavy cells). D) Volcano plot graph of streptavidin-enriched protein SILAC ratios and BH-corrected *P*-values for both SNAP-KEAP1 (red) and KEAP1-SNAP (blue) expressing cells treated with PP1 probe, irradiated and enriched prior to on-bead trypsinolysis and LC-MS/MS analysis (heavy cells). Only the marker proteins KEAP1, MGMT (SNAP-Tag) and PGAM5 are labeled.

PP1 treatment had no significant effect on global protein abundance relative to vehicle treatment (Fig. 3C). Both KEAP1 and SNAP-Tag (a modified MGMT sequence manually included in the search database) proteins were detected in these profiles, and neither showed any differential enrichment in bulk light or heavy proteome under PhotoPPI profiling conditions (Fig. 3C). In stark contrast, the biotin-enriched proteome profile was heavily skewed toward the “heavy”, PP1 probe-treated condition (Fig. 3D). We applied a conservative enrichment cutoff that required a fold-change > 2 and a multiple hypothesis test corrected *P-*value < 0.05 from replicate technical and biological runs to identify proteins that were significantly enriched by PP1 photoproximity profiling. Both KEAP1 and MGMT proteins were at the top of the enriched profile from cells expressing either the C- or N-terminal KEAP1 fusions, confirming the proximal labeling and enrichment of the bait fusion protein (Fig. 3D). PP1 treatment and PhotoPPI processing of cells expressing FLAG-KEAP1 instead of a SNAP-fused KEAP1 construct showed no significant enrichment of any proteins (Suppl. Fig. 6; Suppl. Tables 1-2). The significant bait protein labeling likely results from both within-sphere self-labeling, as well as labeling of adjacent KEAP1 bound in the non-covalent^22^ and covalent^23^ homodimers that are known to form in cells. Indeed, higher enrichment ratios were observed for KEAP1 compared to MGMT, consistent with significant labeling of endogenous KEAP1 bound to the PP1-labeled SNAP-KEAP1 fusions in cells (Fig. 3D). Conspicuously, the next most enriched protein in both profiles was phosphoglycerate mutase 5 (PGAM5), which is a validated KEAP1-interacting protein that harbors a consensus ‘ESGE’ KEAP1 binding site and has been implicated in tethering KEAP1 to the mitochondrial membrane^24-25^. The global PhotoPPI profile also included significant enrichment of proteins involved in vesicle and membrane trafficking, ribosomal biogenesis, mitochondrial membrane transport, splicing, redox regulation, and other functional categories (Suppl. Fig. 7A, C). Like PGAM5, several enriched proteins are known to be localized to the mitochondrial membrane, suggesting that they may interact directly or indirectly with KEAP1. Co-immunoprecipitation experiments with hexokinase 2 (HK2), which was significantly enriched by PP1 but not previously reported as a KEAP1 interactor, confirmed that pulldown of FLAG-KEAP1, but not FLAG-SNAP, results in co-precipitation of HK2 (Suppl. Fig. 7B). HK2 is known to associate with the mitochondrial membrane surface where it can respond to acute changes in ATP levels^26^, suggesting that co-localization of KEAP1 and HK2 at the mitrochondria could enact a dual metabolic and redox sensing complex. Intriguingly, we did not detect NRF2 or CUL3, the transcription factor substrate and ubiquitin-ligase proteins known to associate with KEAP1, in this steady state experiment. Likewise, the only published AP-MS study of KEAP1 binding proteins under basal conditions failed to detect NRF2 or CUL3^27^. Despite completely different proteomic workflows, however, this AP-MS profile had significant overlap with that identified here, with PGAM5 being identified as one of the most enriched proteins in both profiles, as well as similar network-level enrichment of proteins involved in ribosome biogenesis, membrane trafficking, mRNA splicing and detoxification pathways.

### PhotoPPI Detects Dynamic Protein Interaction Rewiring In Cells

To ascertain whether PhotoPPI profiling is capable of capturing dynamic protein interactions we next profiled KEAP1 in cells under conditions of perturbed metabolic or redox states known to result in chemical modification and activation of KEAP1. SILAC-labeled cells expressing the C- or N-terminal SNAP-KEAP1 fusions were treated with either CBR-470-1, a recently described inhibitor of PGK1 that results in modification of KEAP1 by elevated methylglyoxal^23^, or the redox active agent tert-butyl hydroperoxide (tBuOOH). Since both of these conditions are known to activate the KEAP1-NRF2 pathway, we wondered whether distinct interaction profiles could be detected by PhotoPPI. After treatment of both heavy and light cultures with either pharmacologic agent, heavy cells were labeled with the photoprobe and proteomes were processed for quantitative LC-MS/MS as in the basal experiment. Several interactors were clearly retained under both stressed conditions, including the bait proteins KEAP1 and MGMT, as well as PGAM 5 (Fig. 4A-B; Suppl. Tables 3-4). Inspection of the global enrichment profile confirmed profound differences in KEAP1 binding partners in CBR-470-1- and tBuOOH-treated cells compared to the basal profile, as well as many similarities between the two perturbed conditions (Fig. 4C; Suppl. Fig. 7C-E). For example, the trafficking protein SQSTM1, which is implicated in binding to modified KEAP1 and trafficking to authophagic or lysosomal compartments, was highly enriched in the CBR-470-1 treated cells (Suppl. Fig. 7C; Suppl. Table 3). Similarly, both CBR-470-1 and tBuOOH profiles show strong enrichment of lysosomal proteases like CTSB, LGMN and SCPEP1. Other intriguing targets include the transporters like ABCC1, which is a direct transcriptional target of NRF2^28^. A comparative analysis of between PhotoPPI profiles enables drafting of dynamic interaction maps, including KEAP1 interactors that are maintained across conditions (e.g. KEAP1 and PGAM5), present under basal conditions but lost in response to stress (e.g. HK2 and PLK1), or selectively induced in response to stress and KEAP1 modification (e.g. CTSD and LGMN; Fig. 4C). Together this profile enables differential detection of both altered sub-cellular localization and protien-complex interaction for KEAP1 in live cells under basal and stressed conditions (Fig. 4D). These data provide a valuable starting point for future studies into the regulatory roles these interactions plan in KEAP1/NRF2 signaling^29^, and also validate the utility of the PhotoPPI platform to map dynamic interaction profiles in cells.

**Figure 4.**
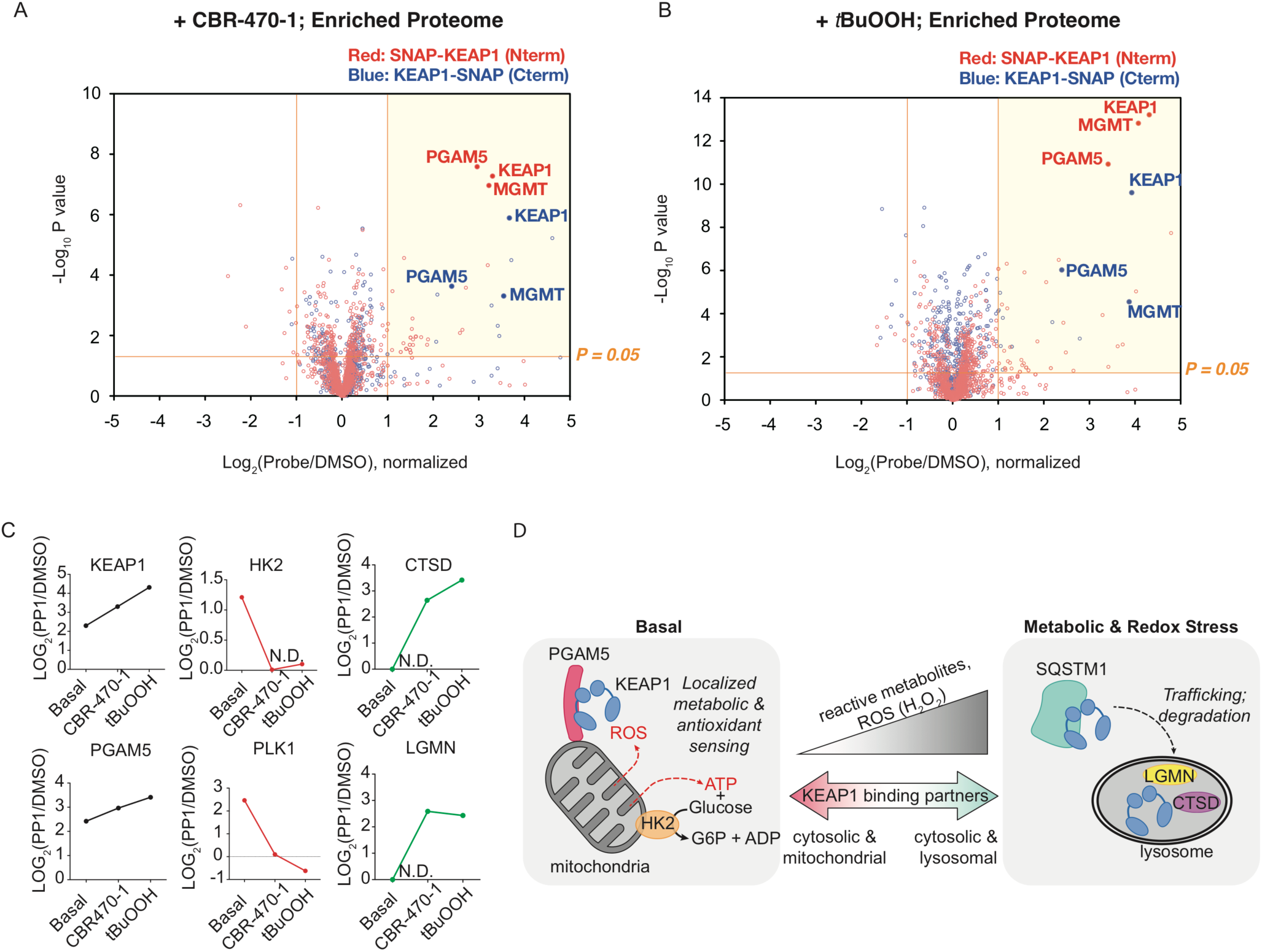
PhotoPPI detects differential KEAP1 localization and protein interactors in response to metabolic and redox stress. A-B) Volcano plots graph of streptavidin-enriched protein SILAC ratios and *P*-values for both SNAP-KEAP1 (red) and KEAP1-SNAP (blue) expressing cells treated with CBR-470-1 (A) or tertbutylhydroperoxide (tBuOOH, B), followed by treatment of heavy cells with PP1 probe, irradiation of both heavy and light and proteomic processing as above in Fig. 3. Only the marker proteins KEAP1, MGMT (SNAP-Tag) and PGAM5 are labeled. C) Graphical depiction of specific target proteins that were significantly enriched in KEAP1 PhotoPPI profiles. Shown are mean enrichment ratios (y-axis) for each protein across all of the runs in each biological condition (x-axis). Proteins that were enriched as KEAP1 interactors among all conditions are labeled in black; proteins that were enriched in the basal profile but lost in stressed conditions are labeled red; proteins that were selectively enriched only in stressed conditions are labeled in green. When a protein was not detected among any runs in a specific condition, a SILAC ratio of 1 (i.e. no enrichment) was assigned and the point labeled as not detected (N.D.). D) Schematic depicting the altered sub-cellular localization and specific protein interactors detected under basal (left) and metabolic/redox stressed conditions (right).

## CONCLUSION

Here we have developed a novel photoproximity protein profiling platform, which relies on complementary photo-responsive chemical probes and genetically encoded SNAP-POI targeting in cells for light-triggered proximity labeling. *In vitro* experiments validated protein-protein interaction-dependent, and light-mediated covalent tagging of proteins bound to a PP1-labeled SNAP-Tag protein. Furthermore, we demonstrated intracellular labeling, light-triggered activation and proteome-wide proximity profiling of KEAP1 in live cells. These studies identified both known and unknown interactions under basal and stress activated conditions. Overall, we believe there are several advantages to the PhotoPPI method that make it attractive for use in specific applications. First, the capacity to load a SNAP-Tag fusion construct with a masked proximity labeling molecule in live cells without significant perturbation to cellular physiology should provide significant advantages relative to other live-cell proximity labeling technologies. For example, the profound rewiring of the KEAP1 interaction profile to redox perturbations like peroxide challenge suggest that employing proximity labeling methods that could perturb the redox environment might miss or mis-assign basal and dynamic interaction states for a protein like KEAP1. Moreover, since the role of redox regulation on protein structure and function is relatively poorly characterized, we cannot be sure which proteins would or would not be affected by such perturbations. Additionally, the facile labeling and washout of free PP1, followed by light-triggered proximity labeling should permit very high temporal and contextual control of the cellular conditions that can be interrogated using the PhotoPPI platform relative to other approaches with prolonged incubation periods or no on/off trigger. We demonstrated the importance of this attribute here by capturing the altered interaction profile of KEAP1 within a time frame shorter than the incubation period for some proximity labeling methods. Indeed, the photocleavage kinetics of even the first generation probe suggest we should be able to probe dynamic events that take place across minutes-long timescales. While not explored here, the light activated control of this system could also allow for selective spatial activation, and therefore proximity profiling, of a unique cellular field. Additionally, an important design element for a high fidelity spatial profiling platform is the reactivity of the tagging group, which sets the practical labeling radius as well as the reactivity profile with target biomolecules. In this regard, the masked carbene nucleophile in our first generation PhotoPPI probe should provide a highly restricted labeling radius, and broader chemical targeting capacity on proximal proteins relative to the acyl phosphate and phenoxyl radicals. We anticipate that guided alteration of the central photocleavage group coupled with tuning of the photoreactive element will enable development of PhotoPPI probes with differential labeling radii and altered target compatibility to provide higher resolution interactome maps. Thus, the PhotoPPI platform could be used alongside existing proximity profiling methods to provide complementary, layered datasets within the same biological contexts. We anticipate that current and future versions of the PhotoPPI platform will be broadly useful in elucidating molecular interaction networks inside living systems.

## ASSOCIATED CONTENT

**Supporting Information. Supporting Information**.

## ACKNOWLEDGMENT

Supplementary Figures 1-7; Supplementary Schemes. Supplementary Tables 1-4.

## AUTHOR INFORMATION

We thank M. Rosner for discussion surrounding the manuscript. We are grateful for financial support of this work from the following: NIH 2T32GM008720-16 (to D.C.M.); NIH R00CA175399 and DP2GM128199-01 (R.E.M.), the Duchoissos Family Institute.

