## Supplemental Information for "Photoproximity Profiling of Protein-Protein Interactions in Cells"

### SUPPORTING MATERIAL FOR
### Photoproximity Profiling of Protein-Protein Interactions in Cells

David C. McCutcheon[1,2‡], Gihoon Lee[1,2‡], Anthony Carlos[1,2], Jeffrey E. Montgomery[1,2], & Raymond E. Moellering[1,2*]

[1]Department of Chemistry & [2]Institute for Genomics and Systems Biology, The University of Chicago; 929 E. 57th Street, Chicago, IL, 60637.

Abbreviations:
benzyl guanine (BnG); photo-FITC benzyl guanine (PF-BnG); [dimethylamino(triazolo[4,5-b]pyridin-3-yloxy)methylidene]-dimethylazanium;hexafluorophosphate (HATU); *N*-hydroxysuccinimide (NHS), *N,N′*-disuccinimidyl carbonate (DSC); 3-(ethyliminomethylideneamino)-*N,N*-dimethylpropan-1-amine;hydrochloride (EDC•HCl); liquid chromatography-mass spectroscopy (LC-MS)

**PF-BnG**

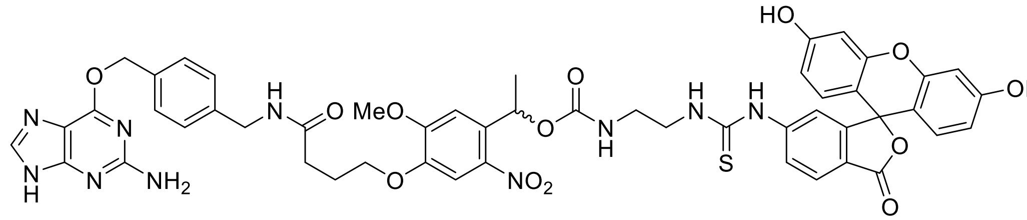

**PP1**

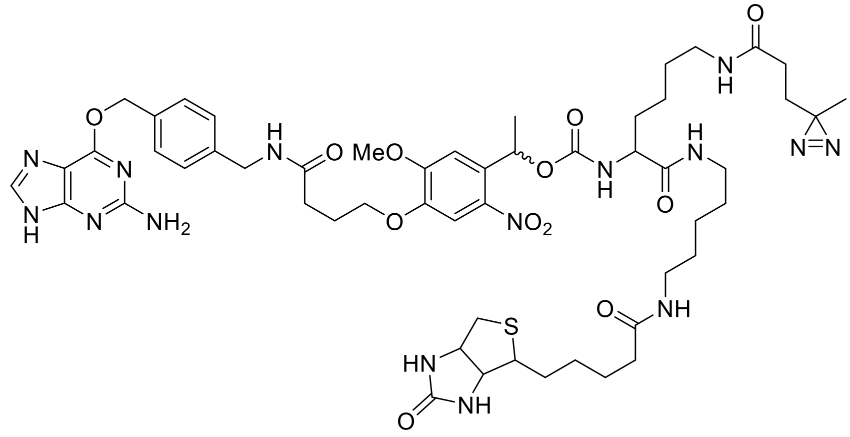

**PP2**

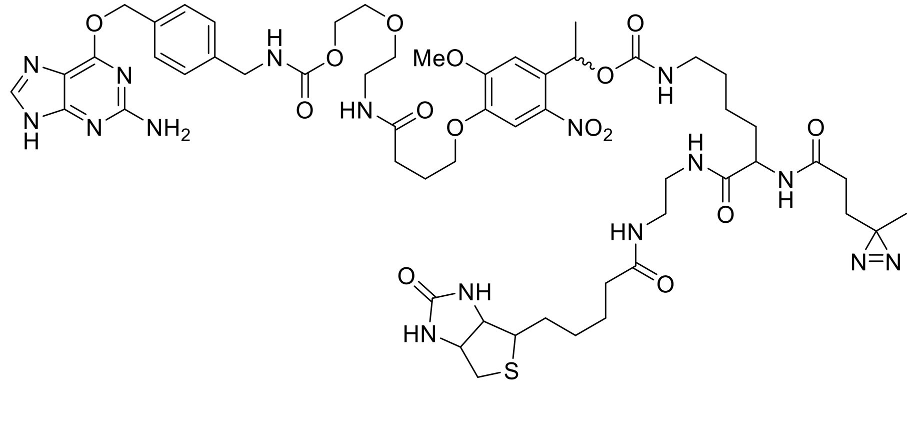

**Supplemental Figure 1:** Chemical structures of photoproximity probes used in this study.

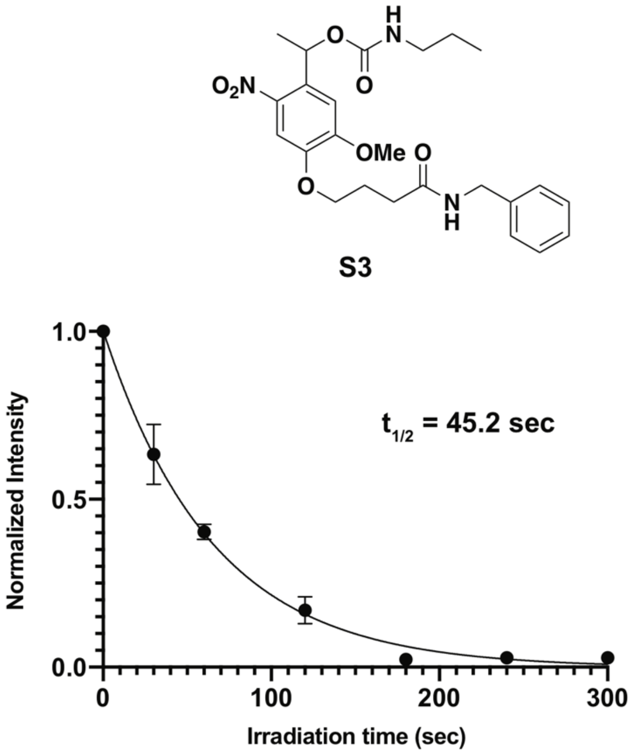


**Supplemental Figure 2:** In vitro photocleavage kinetics. Chemical structures of model photoreactive compounds **S3** and **15** (top)**.** Normalized compound level as measured by LC-MS integration, fit to a one-phase exponential decay model. All data are mean +/- s.e.m. from three technical replicates from a representative biological replicate. Curve fit in Prism 8.

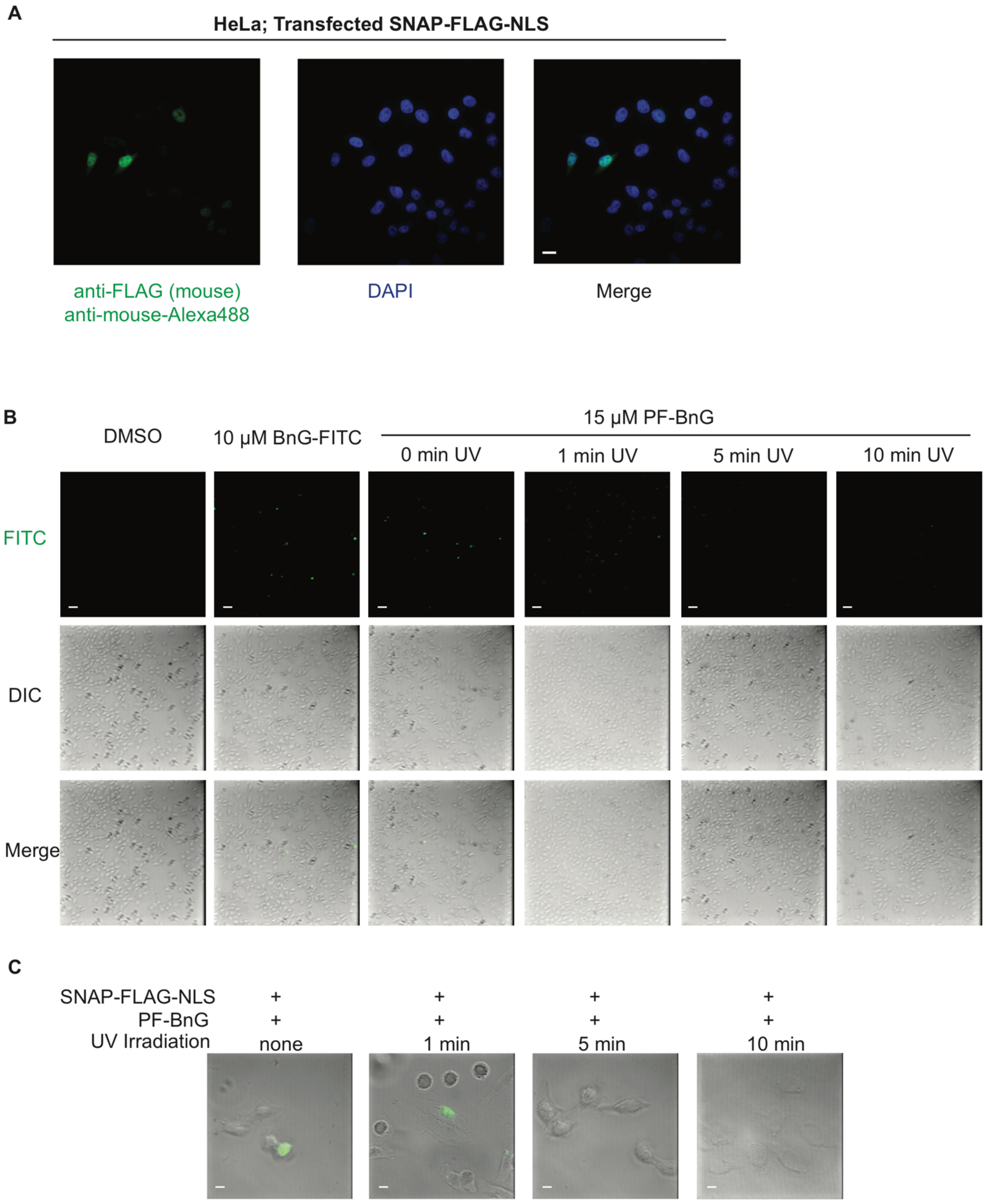


**Supplemental Figure 3:** Sub-cellular localization and photocleavage with the PhotoPPI system. **A**) Fluorescence microscopy images of HeLa cells transiently transfected with SNAP-FLAG-NLS, which exhibit strong nuclear localization of the SNAP protein. **B**) Fluorescence and brightfield microscopy images of HeLa cells transiently-transfected with SNAP-FLAG-NLS show nuclear-localization of FITC signal in cells treated with the non-cleavable

FITC-BnG, or PF-BnG in the absence of UV irradiation (left three columns). Progressive exposure to UV-irradiation at 365 nm results in complete loss of FITC signal in less than 10 min. Note that DAPI co-staining of the nucleus could not be performed due to overlap in the wavelength for DAPI excitation and nitroveratryl cleavage. **C**) Magnified view of cells in (B) above and reproduced from Fig. 1C here for comparison. Scale bars in A, B and C equal 8.3, 50 and 8.3 μm, respectively.

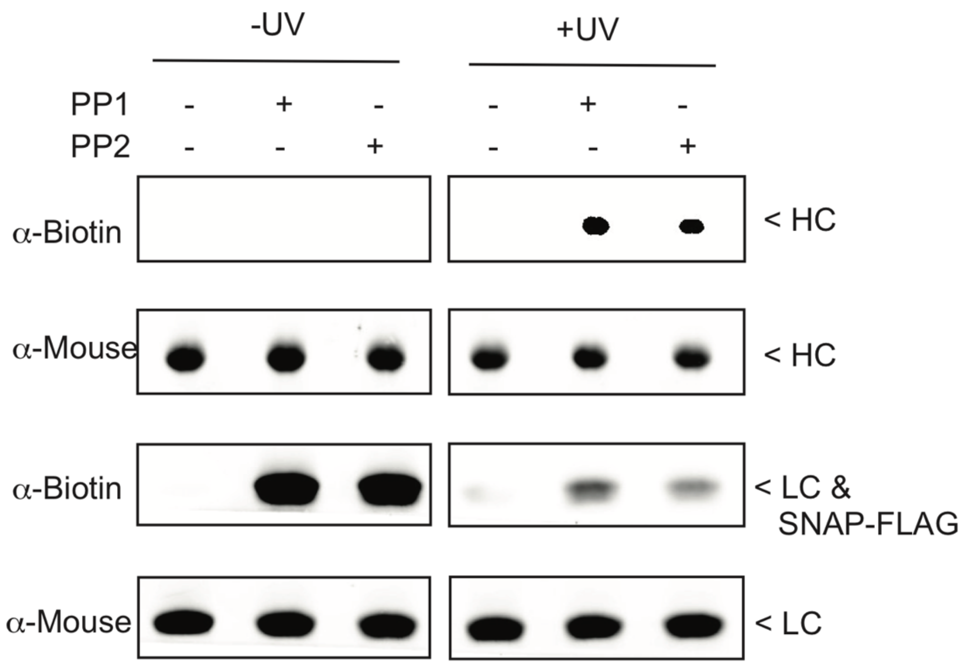


**Supplemental Figure 4:** In vitro photoproximity labeling in the presence of whole cell proteome. Anti-biotin (streptavidin-800) and anti-mouse Western blot analyses of PP1 labeled SNAP-FLAG/ α-FLAG antibody complex with and without UV irradiation prior to analysis. Photolabeling was performed in the presence of whole cell lysate. Labels for individual proteins are included at appropriate molecular weights: LC, light chain; HC, heavy chain; "SNAP" label represents SNAP-Tag protein without the FLAG epitope.

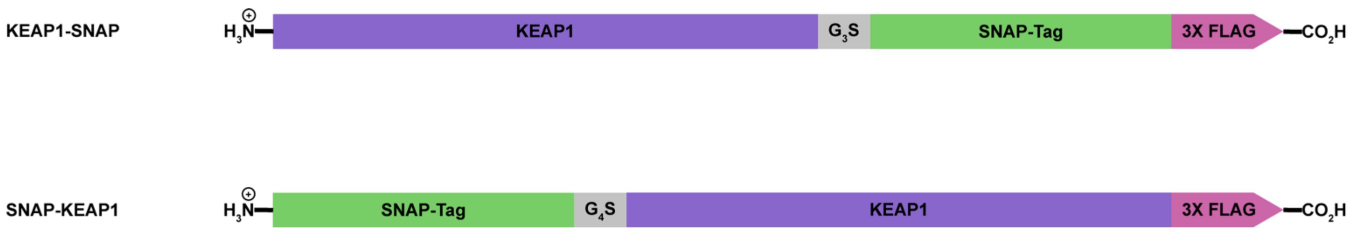


**Supplemental Figure 5:** Schematic depicting C-terminal (KEAP1-SNAP) and N-terminal (SNAP-KEAP1) genetic fusions used in photoproximity profiling of KEAP1 in cells. $G_xS$ represents a glycine-serine spacer, with X indicating the number of glycines.

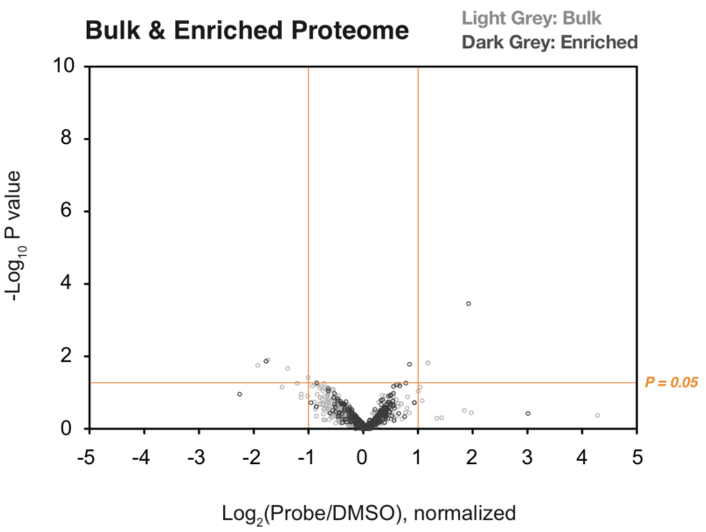


**Supplemental Figure 6:** Overlaid volcano plots of bulk and biotin-enriched proteomic runs from SILAC cells expressing FLAG-KEAP1, treated with DMSO (light) or PP1 (heavy), irradiated with 365 nm light, and processed according to the PhotoPPI workflow. In contrast to cells expressing SNAP-KEAP1 constructs (Fig. 3 & 4), these mock cells show no enrichment.

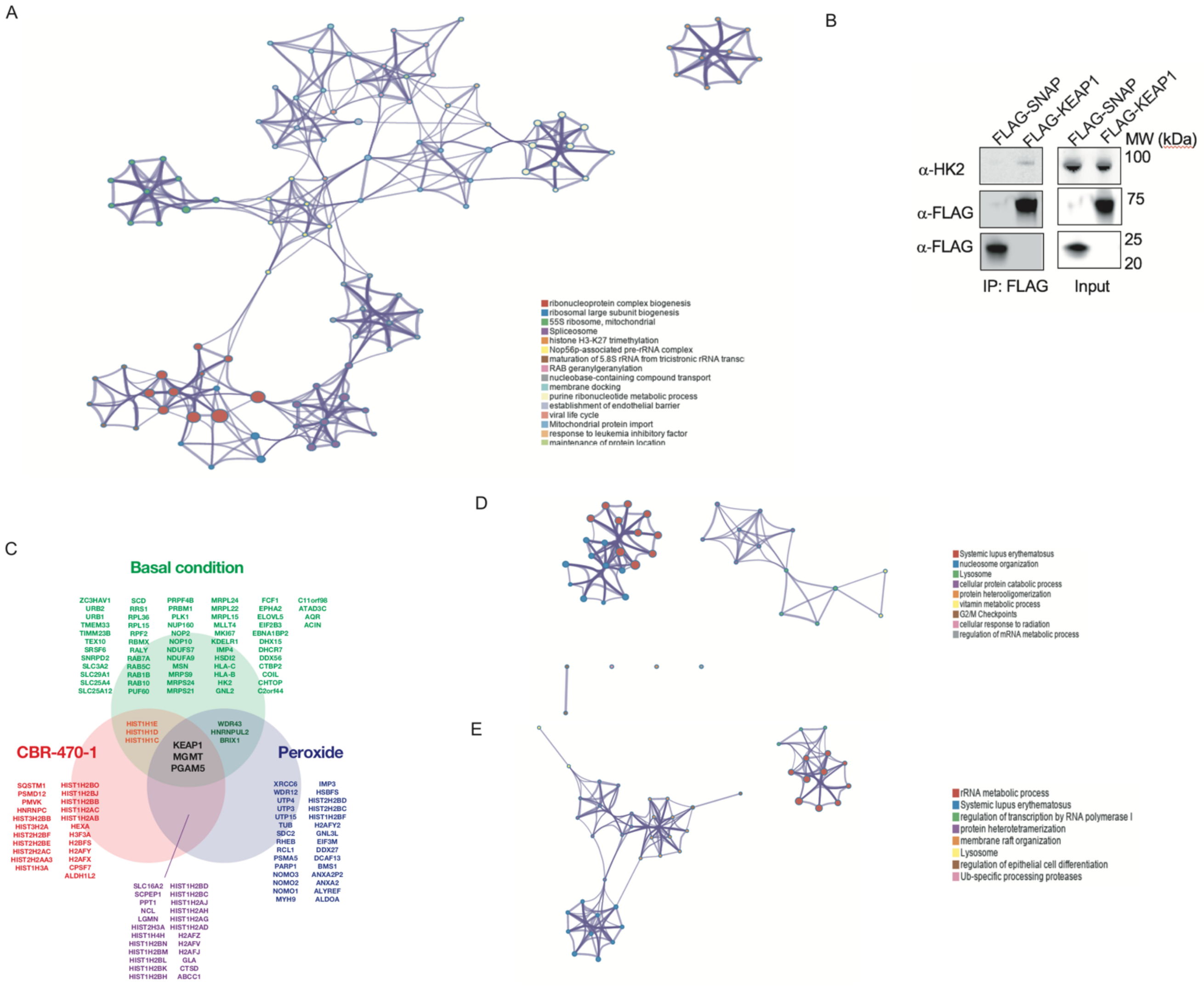


**Supplemental Figure 7.** A) Metascape network analysis of significantly enriched protein functional groups and ontology categories in the basal KEAP1 PhotoPPI-enriched profile. B) Co-immunoprecipitation of FLAG-SNAP and FLAG-KEAP1 from stably expressing HeLa cells. Western blot analysis of FLAG-SNAP, FLAG-KEAP1 and HK2 proteins demonstrate selective enrichment of HK2 by FLAG-KEAP1 but not FLAG-SNAP. C) A) Venn-diagram depicting enriched proteins under basal, CBR-470-1 and tBuOOH treatment conditions. Proteins with non-unique peptides that may be redundant between isoforms are included in this list. D-E) network analysis of significantly enriched protein functional groups and ontology categories in the CBR-470-1 (D) and tBuOOH (E) KEAP1 PhotoPPI-enriched profiles. Redundant protein identifications due to non-unique peptides were merged in this analysis to avoid overrepresentation.

**General synthetic methods**
Reagents purchased from commercial suppliers were analytical grade and used without further purification. All reactions were carried out in oven dried flasks using anhydrous solvents (Acros) unless otherwise specified. Reaction progress was monitored by thin-layer chromatography on Macherey-Nagel SIL G-25 UV254 TLC plates, visualized with UV light, ceric ammonium molybdate (CAM), *p*-anisidine, bromophenol blue, 2,4-dinitrophenyl hydrazine (DNP), or KMnO4 TLC stains. Nuclear magnetic resonance spectra were acquired using either a Bruker AVANCE II+ 500; 11.7 Tesla NMR or Bruker DRX 400; 9.3 Tesla NMR instrument. Accurate mass measurements were obtained using an Agilent 6224 Tof-MS instrument. When necessary, compounds were purified via flash column chromatography using Siliaflash F60 60 Å, 230-400 mesh silica gel (Silicycle)

**Chemical synthesis of photoproximity profiling (PhotoPPI)-probes**

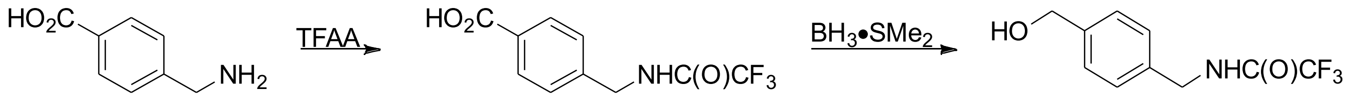


1 2 3

**4-[[(2,2,2-trifluoroacetyl)amino]methyl]benzoic acid (2)**
Solid 4-(aminomethyl)benzoic acid **1** (15.1 g, 100 mmol) was dissolved in TFAA (42 mL) cooled to 0 °C. Once dissolved, the ice bath was removed and the reaction was allowed to stir at rt until starting material was consumed, ~2 hr. Upon completion, the reaction was quenched with $H_2O$ (100 mL) and precipitate collected via vacuum filtration. The product was dried under suction then collected to afford benzoic acid **2** (24.0 g 97%) as a white solid.
$^1$H NMR (500 MHz, DMSO-d$^6$) $\delta$ 7.93 (d, $J$ = 8.3 Hz, 2H), 7.39 (d, $J$ = 8.3 Hz, 2H), 4.47 (d, J = 6.0 Hz, 2H). $^{13}$C NMR (125 MHz, DMSO-d$^6$) $\delta$ 167.12, 156.58 (q, $J$ = 36.3 Hz), 142.50, 129.89, 129.65, 127.44, 116.05 (q, J = 228.1 Hz), 42.41.

**2,2,2-trifluoro-*N*-[[4-(hydroxymethyl)phenyl]methyl]acetamide (3)**
Borane dimethylsulfide (13.8 mL, 145 mmol) was added dropwise to an anhydrous THF (483 mL) solution of benzoic acid **2** (11.9 g, 48.3 mmol) while maintaining an internal temperature of 0 °C. After complete addition the ice bath was removed and the mixture was stirred at rt overnight. The reaction was quenched with MeOH (100 mL) and stirred at rt for an additional 1 hr. The volatiles were removed and the residue taken up in EtOAc. Impurities were removed by successive washes with 1M NaOH, $H_2O$, and brine. The organics were then dried over $Na_2SO_4$, filtered, and concentrated *in vacuo*. Subsequent column chromatographic purification of the residue, eluting with 20:1 ($CH_2Cl_2$:MeOH), provided acetamide **3** (10.4 g, 92%) as a white solid.
$^1$H NMR (500 MHz, $CDCl_3$-d) $\delta$ 7.35 (d, $J$ = 8.2 Hz, 1H), 7.27 (d, $J$ = 8.0 Hz, 1H), 4.67 (s, 1H), 4.50 (d, $J$ = 5.8 Hz, 1H). $^{13}$C NMR (125 MHz, CDCl3-d) $\delta$ 157.32 (q, $J$ = 36.7 Hz), 141.11, 135.33, 128.29, 127.67, 115.97 (q, $J$ = 287.8 Hz), 64.90, 43.75.

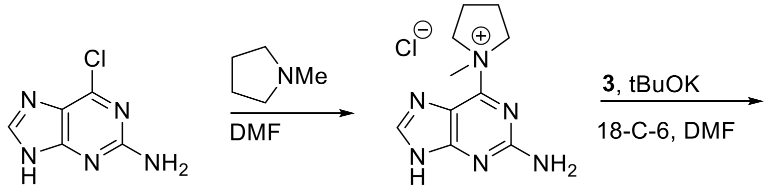


**4** **5**

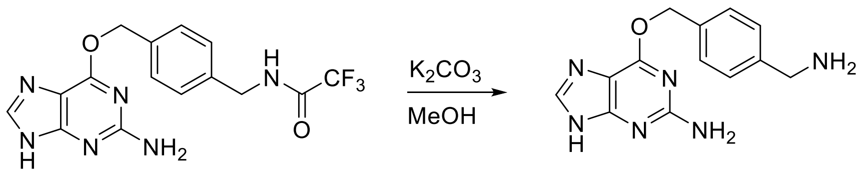


**6** **7**

**6-(1-methylpyrrolidin-1-ium-1-yl)-7H-purin-2-amine chloride (5)**

Neat *N*-methylpyrollidine (7.80 mL, 73.7 mmol) was added to an anhydrous DMF (144 mL) solution of 6-chloro-7H-purin-2-amine **4** (5.00 g, 29.5 mmol) and stirred at 40 °C overnight. The resultant chloride salt **5** (5.45 g, 73%) was collected via vacuum filtration, dried under suction, and used without further purification.

***N*-[[4-[(2-amino-7H-purin-6-yl)oxymethyl]phenyl]methyl]-2,2,2-trifluoroacetamide (6)**

An oven dried round-bottom flask containing methylpyrrolidinium chloride **4** (6.90 g, 27.1 mmol), acetamide **3** (12.6 g, 54.2 mmol), potassium tert-butoxide (12.2 g, 108 mmol), and 18-c-6 (1.07 g, 4.07 mmol) in 54 mL DMF was stirred for 6 hrs at 50 °C. Upon completion the solvent was evaporated and the crude residue absorbed to silica. Purification was achieved using column chromatography eluting with a gradient of MeOH in $CH_2Cl_2$ (2 – 10%) to give the trifluoroacetamide protected amine **6** (8.94 g, 90%) as a white solid.

$^{1}$H NMR (500 MHz, DMSO-$d^6$) $\delta$ 7.81 (s, 1H), 7.49 (d, $J$ = 7.9 Hz, 2H), 7.30 (d, $J$ = 8.0 Hz, 2H), 5.46 (s, 2H), 4.4 (d, $J$ = 5.9 Hz, 2H).

**6-[[4-(aminomethyl)phenyl]methoxy]-7H-purin-2-amine (7)**

Trifluoroacetamide **6** (2.59 g, 7.06 mmol) was added to a suspension of $K_2CO_3$ (4.84 g, 35.0 mmol) in 21 mL of MeOH:$H_2O$ (20:1) and stirred vigorously overnight at 50 °C. Upon consumption of the starting material the mixture was filtered through a pad of celite, washing with MeOH. The filtrate was concentrated *in vacuo* and the residue taken up in 10 mL $H_2O$. While cooling, the pH was adjusted to ~7 with HCl. The resultant precipitate was isolated via suction filtration and washed with cold water to yield amine **7** (1.77 g, 93%) as a white solid. $^{1}$H NMR (500 MHz, DMSO-$d^6$) $\delta$ 7.82 (s, 1H), 7.43 (d, $J$ = 7.9 Hz, 2H), 7.34 (d, $J$ = 7.8 Hz, 2H), 5.45 (s, 2H), 3.71 (s, 2H). $^{13}$C NMR (125 MHz, DMSO-$d^6$) $\delta$ 159.65, 157.73, 143.72, 140.45, 135.00, 134.67, 128.47, 127.14, 126.98, 66.66, 45.27.

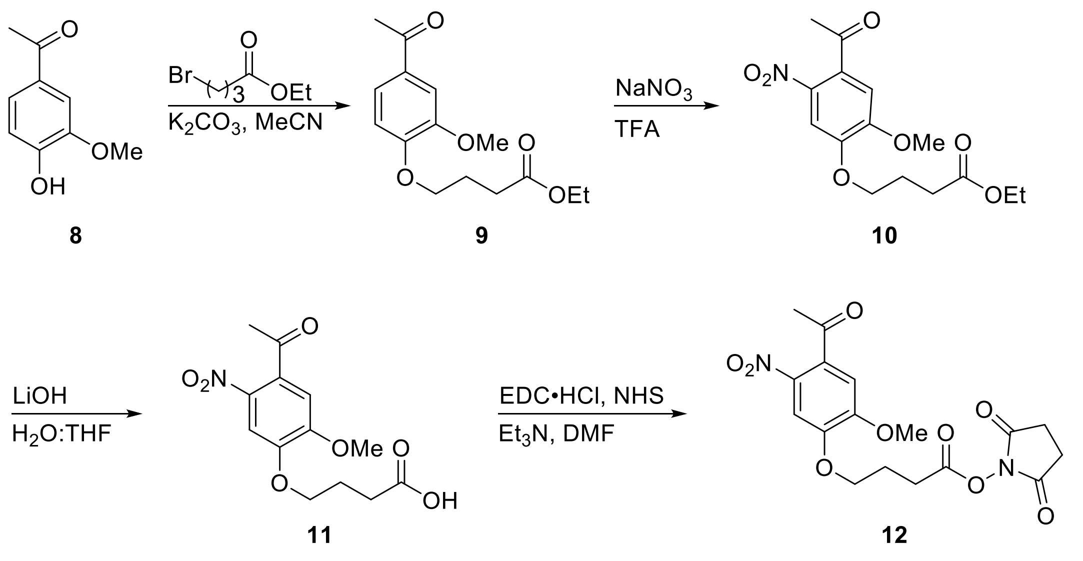

**ethyl 4-(4-acetyl-2-methoxyphenoxy)butanoate (9)**
Solid 1-(4-hydroxy-3-methoxyphenyl)ethanone **8** (8.31 g, 50 mmol) was added to a suspension of $K_2CO_3$ (69.1 g, 500 mmol) in 100 mL of anhydrous MeCN. Ethyl 4-bromobutanoate (14.3 mL, 100 mmol) was added and the mixture stirred overnight at 60 °C. Upon consumption of starting material the mixture was filtered over a pad of celite, washing with cold MeCN. Volatiles were evaporated and the residue recrystallized from $Et_2O$ to provide ester **9** (13.2 g, 94%) as a white powder. $^1$H NMR (400 MHz, CDCl3-*d*) δ 7.57 (dd, *J* = 8.3, 2.1 Hz, 1H), 7.55 (d, *J* = 2.0 Hz, 1H), 6.92 (d, *J* = 8.3 Hz, 1H), 4.21 – 4.13 (m, 4H), 3.94 (s, 3H), 2.59 (s, 3H), 2.56 (t, *J* = 7.2 Hz, 2H), 2.26 – 2.17 (m, 2H), 1.28 (t, *J* = 7.1 Hz, 3H).

**ethyl 4-(4-acetyl-2-methoxy-5-nitrophenoxy)butanoate (10)**
Using an ice bath, 50 mL of trifluoroacetic acid (50 mL) was cooled to 0 °C prior to the addition of ester **9** (12.8 g, 45.7 mmol). Solid $NaNO_3$ (11.6 g, 136 mmol) was added in portions to the stirred mixture maintaining 0 °C. Upon completion the reaction was quenched with 200 mL of $H_2O$, the resulting precipitate filtered, then dried under suction to afford the nitro arene **10** (13.8 g, 93%) as a yellow powder. $^1$H NMR (400 MHz, CDCl3) *δ* 7.61 (s, 1H), 6.74 (s, 1H), 4.21 – 4.12 (m, 4H), 3.95 (s, 3H), 2.54 (t, *J* = 7.2 Hz, 2H), 2.49 (s, 3H), 2.25 – 2.16 (m, 2H), 1.27 (t, *J* = 7.2 Hz, 3H).

**4-(4-acetyl-2-methoxy-5-nitrophenoxy)butanoic acid (11)**
A solution of nitro arene **10** (2.52 g, 7.76 mmol) in THF (76 mL) was combined with 39 mL of 2M LiOH. The reaction was allowed to stir vigorously for 3 hrs at rt. Upon starting material consumption, the reaction was quenched with 1M $NaHSO_4$ (100 mL) and extracted with EtOAc. The combined organic extracts were dried over $NaSO_4$ and filtered. Precipitate that formed upon solvent removal *in vacuo* was filtered and dried under suction to provide the title acid **11** (1.86 g, 81%) as a pale yellow powder. $^1$H NMR (400 MHz, $CDCl_3$) *δ* 7.62 (s, 1H), 6.75 (s, 1H), 4.17 (t, *J* = 6.2 Hz, 2H), 3.95 (s, 3H), 2.63 (t, *J* = 7.1 Hz, 2H), 2.49 (s, 3H), 2.27 – 2.13 (m, 2H).

**(2,5-dioxopyrrolidin-1-yl) 4-(4-acetyl-2-methoxyphenoxy)butanoate (12)**
Acid **11** (4.65 g, 15.6 mmol) was added to an anhydrous DMF (30 mL) solution containing EDC•HCl (4.48 g, 23.4 mmol) and NHS (2.69 g, 23.4 mmol). The mixture was stirred overnight at rt. Addition of chilled $Et_2O$ to the bulk solution resulted in precipitate, which was collected via vacuum filtration and dried under suction to yield NHS-ester **12** (5.47 g, 89%) as a yellow powder. $^1$H NMR (500 MHz, $CDCl_3$) *δ* 7.63 (s, 1H), 6.75 (s, 1H), 4.21 (t, *J* = 6.0 Hz, 2H), 3.96 (s,

3H), 2.89 (t, $J$ = 7.3 Hz, 2H), 2.85 (bs, 4H), 2.49 (s, 3H), 2.35 – 2.28 (m, 2H). $^{13}$C NMR (125 MHz, $CDCl_3$) $\delta$ 200.28, 169.20, 168.18, 154.54, 148.71, 133.29, 108.88, 108.44, 67.66, 56.72, 30.58, 27.62, 25.73, 24.21.

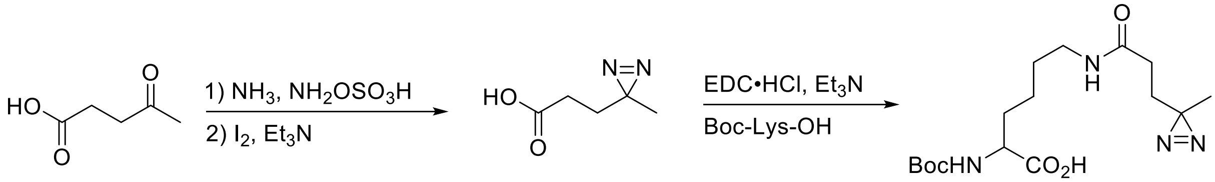


**13** **14** **15**

**3-(3-methyldiazirin-3-yl)propanoic acid (14)**

Gaseous ammonia (*ca.* 200 mL) was condensed at -78 °C into an oven dried two-neck round-bottom flask, 4-oxopentanoic acid **13** (11.6 g, 100 mmol) added, then the mixture refluxed at 0 °C for 4 hrs. A suspension of amino hydrogen sulfate (14.0 g, 123 mmol) in anhydrous MeOH (150 mL) was added via addition funnel over a 45 min period maintaining 0 °C. The heterogenous reaction was then vigorously stirred overnight, allowing the ammonia to slowly evaporate as the temperature rose to rt. The resulting slurry was filtered over a pad of celite, washing the solids with MeOH. The solvent was reduced under vacuum (50 mL) to ensure removal of residual ammonia. The crude residue was then diluted with MeOH (100 mL) and cooled to 0 °C before the addition of $Et_3N$ (20.8 mL, 150 mmol). Solid $I_2$ (25.5 g, 100 mmol) was added in portions until the color of iodine persisted and the reaction stirred for 2 hrs allowing the temperature to rise to rt. Upon completion, the volatiles were removed in vacuo and the residue diluted with EtOAc (150 mL). The organics were successively washed with 1 M NaHSO4 (50 mL x 2), 0.5 M $Na_2S_2O_3$ (50 mL), brine (50 mL) then dried over $NaSO_4$. The combined organics were filtered and absorbed to silica then subjected to chromatographic purification eluting with a gradient of MeOH in $CH_2Cl_2$ (2 – 5%). The title acid **14** (7.29 g, 57%) was isolated as a thin red oil. $^1$H NMR (500 MHz, $CDCl_3$) $\delta$ 11.18 (bs, 1H), 2.18 – 2.14 (m, 2H), 1.68 – 1.60 (m, 2H), 0.95 (s, 3H). $^{13}$C NMR (125 MHz, $CDCl_3$) $\delta$ 178.66, 29.24, 28.48, 25.05, 19.55.

**(2S)-6-[[3-(3-methyldiazirin-3-yl)propanoyl]amino]-2-[(2-methylpropan-2-yl)oxycarbonyl-amino]hexanoic acid (15)**

HATU (910 mg, 2.39 mmol) was added to a 20 mL vial charged with acid **14** (323 mg, 2.52 mmol), *i*$Pr_2NEt$ (1.30 mL, 7.87 mmol) in anhydrous DMF (12 mL). The mixture was stirred for 1 hr at rt after which Boc-Lys-OH (621 mg, 2.52 mmol) was added in one portion and stirring continued overnight. Upon completion, the reaction was diluted with EtOAc and washed successively with 1 M NaHSO4 (20 mL x 2), $H_2O$ (20 mL x 2), and brine (20 mL). The organics were filtered and concentrated before being purified by reverse phase HPLC, eluting with a gradient of MeOH in $H_2O$ (0 – 95%). The desired acid **15** (381 mg, 42%) was obtained in modest yield as a beige solid. $^1$H NMR (400 MHz, DMSO-$d^6$) $\delta$ 12.4 (s, 1H), 7.83 (t, $J$ = 5.6 Hz, 1H), 7.01 (d, $J$ = 8.0 Hz, 1H), 3.81 (ddd, $J$ = 9.5, 7.9, 4.7 Hz, 1H), 3.00 (q, $J$ = 6.4 Hz, 2H), 1.93 (dd, $J$ = 8.5, 6.9 Hz, 2H), 1.67 – 1.59 (m, 1H), 1.59 – 1.52 (m, 4H), 1.37 – 1.27 (m, 12H), 0.97 (s, 3H). $^{13}$C NMR (125 MHz, DMSO-$d^6$) $\delta$ 174.28, 170.50, 155.61, 77.95, 53.43, 38.24, 30.41, 29.88, 29.79, 28.68, 28.23, 25.85, 23.07, 19.34.

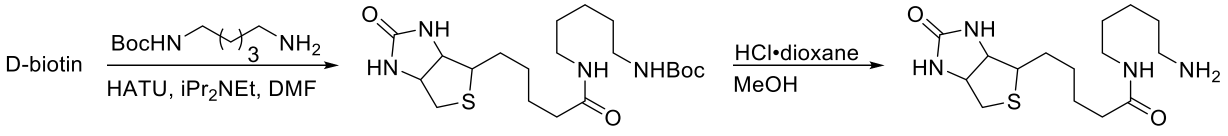


**16** **17**

##### (3aS,4S,6aR)-*N*-[5-[(1,1-dimethylethoxy)carbonyl]aminopentyl]hexahydro-2-oxo-1H-thieno[3,4-d]imidazole-4-pentanamide (16)

A solution of D-biotin (1.95 g, 7.98 mmol), *i*$Pr_2NEt$ (4.20 mL, 24.1 mmol) and HATU (6.08 g, 16.0 mmol) in anhydrous DMF (20 mL) was stirred for 1 hr at rt. Neat tert-butyl *N*-(5-aminopentyl)carbamate (1.78 g, 8.78 mmol) was added and the reaction stirred overnight. Upon completion $Et_2O$ was flowed in and the resultant precipitate filtered under vacuum. The crude solid was purified by iterative recrystallization from acetone and hexanes then $H_2O$ and acetone. Amide **16** (2.95 g, 80%) was isolated by vacuum filtration as a white powder. $^1$H NMR (500 MHz, DMSO-$d^6$) $\delta$ 7.73 (t, *J* = 5.6 Hz, 1H), 6.76 (t, *J* = 5.7 Hz, 1H), 6.43 (s, 1H), 6.36 (s, 1H), 4.30 (dd, *J* = 7.6, 5.0 Hz, 1H), 4.12 (ddd, *J* = 7.7, 4.4, 1.9 Hz, 1H), 3.09 (ddd, *J* = 8.7, 6.1, 4.4 Hz, 1H), 2.99 (q, *J* = 6.6 Hz, 2H), 2.87 (q, *J* = 6.7 Hz, 2H), 2.82 (dd, *J* = 12.4, 5.1 Hz, 1H), 2.57 (d, *J* = 12.4 Hz, 1H), 2.03 (t, *J* = 7.4 Hz, 2H), 1.60 – 1.57 (m, 1H), 1.54 – 1.41 (m, 3H), 1.36 – 1.17 (m, 20H). $^{13}$C NMR (125 MHz, DMSO-$d^6$) $\delta$ 171.78, 162.71, 155.58, 77.32, 61.05, 59.19, 55.46, 38.35, 35.23, 29.20, 28.89, 28.30 (4C), 28.25, 28.06, 25.36, 23.74.

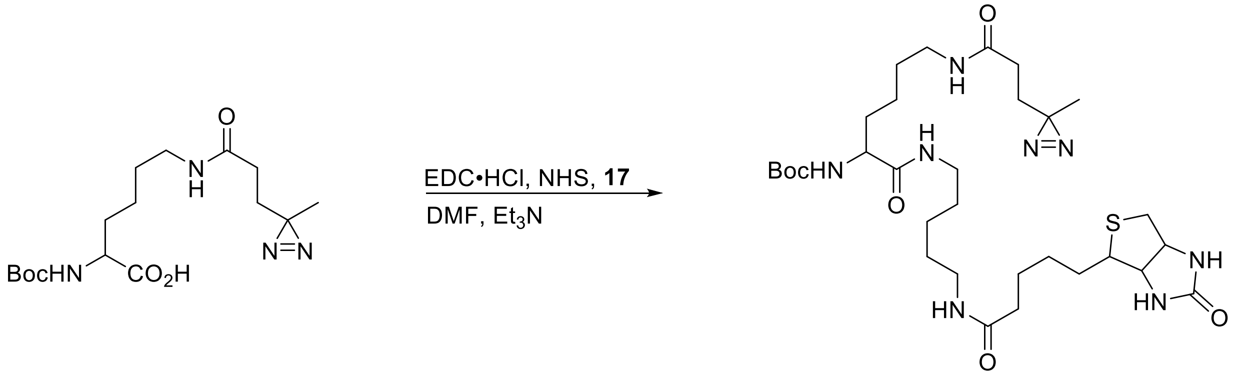


**15** **18**

##### tert-butyl *N*-[6-[[3-(3-methyldiazirin-3-yl)propanoyl]amino]-1-oxo-1-[5-[5-(2-oxo-1,3,3a,4,6,6a-hexahydrothieno[3,4-d]imidazol-4-yl)pentanoylamino]pentylamino]hexan-2-yl]carbamate (18)

Amide **16** (1.80 g, 4.20 mmol) was dissolved with 4 M HCl in 1,4-dioxane (21 mL) and MeOH (1 mL). The mixture stirred at rt for 1 hr after which the volatiles were evaporated. The crude residue was taken up in 20 mL of MeOH and chilled with an ice bath before the addition of 7 M ammonia in MeOH (6 mL). The organics were reduced under vacuum and filtered to remove solids. The residue resulting from filtrate concentration was recrystallized from MeOH in $Et_2O$ to afford amine **17** in quantitative yield and was used without further purification. A separate round-bottom flask containing acid **15** (712 mg, 2.00 mmol), HATU (912 mg, 2.40 mmol), and $Et_3N$ (417 µL, 3.0 mmol) in anhydrous MeCN (10 mL) was stirred at rt for 4 hrs. Afterward, an anhydrous DMSO (7 mL) solution of amine **17** (788 mg, 2.40 mmol) was flowed in and the mixture stirred overnight. Upon completion, the reaction was diluted with EtOAc and washed successively with 1 M $NaHSO_4$ (20 mL), $H_2O$ (20 mL x 2), brine (20 mL) then dried over $NaSO_4$. The crude residue was purified by preparatory reverse phase HPLC, eluting with a gradient of MeOH in $H_2O$ (50 – 95%). Carbamate **18** (453 mg, 34%) was isolated as a waxy solid. $^1$H NMR (500

MHz, $CD_3OD$) $\delta$ 4.50 (dd, $J$ = 7.9, 4.8 Hz, 1H), 4.31 (dd, $J$ = 7.9, 4.4 Hz, 1H), 3.95 (dd, $J$ = 8.9, 5.2 Hz, 1H), 3.24 – 3.13 (m, 7H), 2.93 (dd, $J$ = 12.7, 5.0 Hz, 1H), 2.71 (d, $J$ = 12.7 Hz, 1H), 2.20 (t, $J$ = 7.4 Hz, 2H), 2.07 (dd, $J$ = 8.5, 6.9 Hz, 2H), 1.79 – 1.30 (m, 29H), 1.01 (s, 3H). $^{13}$C NMR (125 MHz, $CD_3OD$) $\delta$ 175.80, 175.03, 174.29, 165.98, 157.68, 80.47, 63.31, 61.54, 56.99, 56.13, 41.05, 40.17, 40.13, 40.06, 36.78, 33.06, 31.49, 31.31, 29.97, 29.93, 29.76, 29.46, 28.74, 26.89, 26.33, 25.12, 24.26, 19.76.

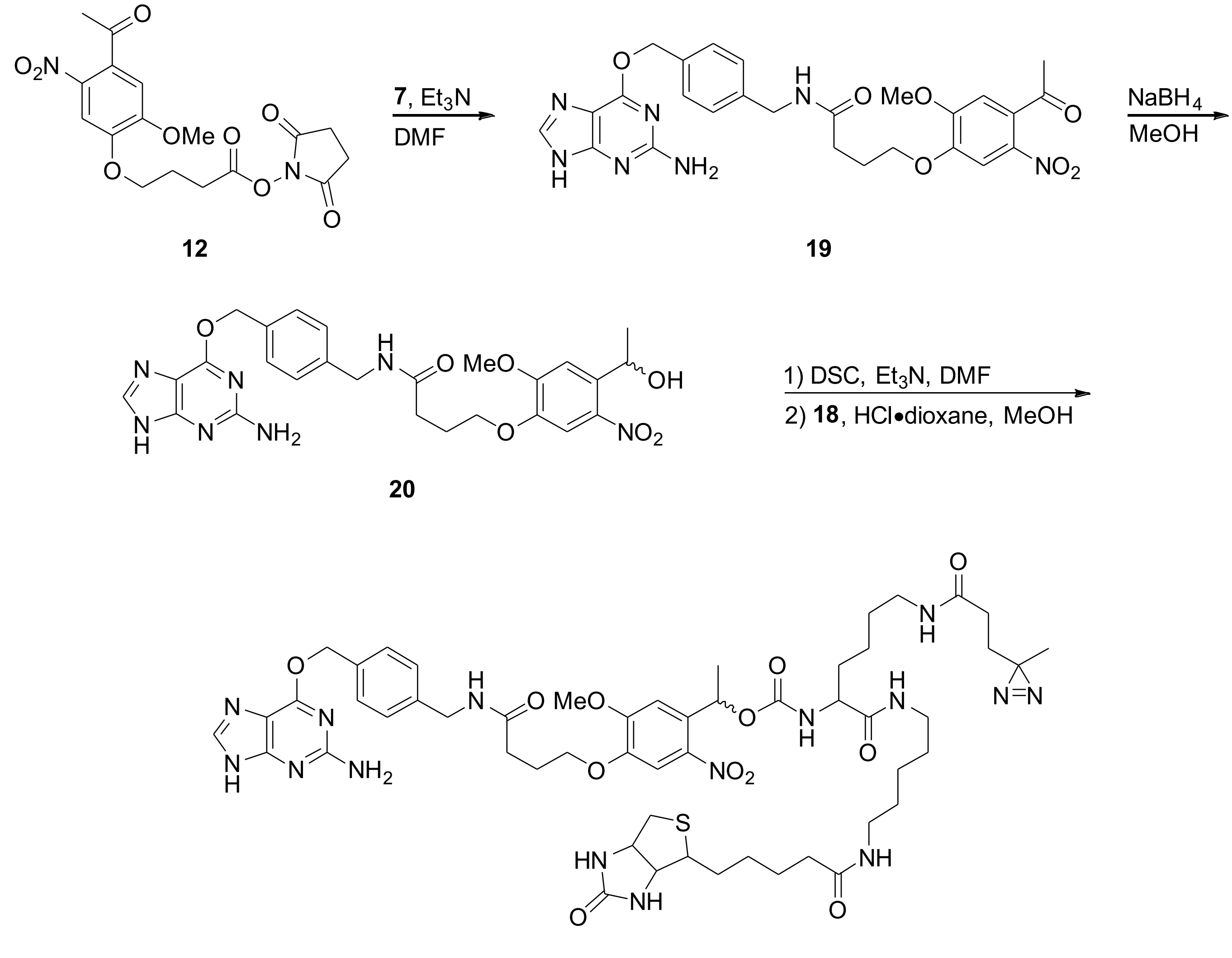


**PP1**

#### Ketone (19)

A DMF (16 mL) solution of NHS-ester **12** (2.30 g, 5.83 mmol) was added to a suspension of amine **7** (1.50 g, 5.55 mmol) and *i*$Pr_2NEt$ (2.75 mL, 16.6 mmol) in anhydrous DMF (25 mL). The resulting homogenous mixture was stirred at rt for 5 hrs. Upon consumption of starting material, chilled $Et_2O$ was flowed into the reaction and supernatant decanted from resultant oil. The crude residue was taken up in $CH_2Cl_2$ and sonicated until formation of precipitate, which was isolated by filtration. The solids were dried under suction to yield pure ketone **19** (1.32 g, 43%) as a beige powder. $^1$H NMR (500 MHz, DMSO-$d^6$) $\delta$ 8.42 (t, $J$ = 5.9 Hz, 1H), 8.19 (s, 1H), 7.62 (s, 1H), 7.46 (d, $J$ = 7.7 Hz, 2H), 7.27 (d, $J$ = 7.8 Hz, 2H), 7.23 (s, 1H), 5.48 (s, 2H), 4.28 (d, $J$ = 5.9 Hz, 2H),

4.12 (t, $J$ = 6.4 Hz, 2H), 3.92 (s, 3H), 2.51 (s, 3H), 2.33 (t, $J$ = 7.4 Hz, 2H), 2.00 (p, $J$ = 6.9 Hz, 2H).
$^{13}$C NMR (125 MHz, DMSO-d$^6$) $\delta$ 199.35, 171.42, 159.13, 158.88, 154.76, 154.19, 153.28, 148.58, 139.83, 138.36, 134.59, 131.11, 128.77, 127.31, 109.84, 107.96, 68.55, 67.38, 56.67, 41.89, 31.47, 30.73, 30.05, 24.58.

**Alcohol (20)**
Solid $NaBH_4$ (113 mg, 3.00 mmol) was added in portions to a rt methanolic solution (10 mL) of ketone **19** (550 mg, 1.00 mmol) with vigorous stirring. Upon completion, the reaction was passed through a plug of silica to remove inorganics. The filtrate was concentrated in vacuo and the residue sonicated in $Et_2O$. The resultant precipitate was isolated via vacuum filtration then dried under suction to yield alcohol **20** (395 mg, 72%) as a yellow powder. $^1$H NMR (400 MHz, $CD_3OD$) $\delta$ 7.84 (bs, 1H), 7.52 (s, 1H), 7.42 (d, $J$ = 7.9 Hz, 2H), 7.37 (s, 1H), 7.26 (d, $J$ = 8.0 Hz, 2H), 5.49 (s, 2H), 5.44 (q, $J$ = 6.2 Hz, 1H), 4.36 (s, 2H), 4.04 (t, $J$ = 6.1 Hz, 2H), 3.91 (s, 3H), 2.46 (t, $J$ = 7.3 Hz, 2H), 2.12 (p, $J$ = 6.6 Hz, 2H), 1.45 (d, $J$ = 6.3 Hz, 3H).

**Photoproximity Probe one (PP1)**
A solution of carbamate **18** (270 mg, 405 μmol) in MeOH (1 mL) was added to 4 M HCl•dioxane (2 mL) at 0 °C. The temperature was allowed to come to rt while stirring for 2 hrs. Upon disappearance of starting material the volatiles were evaporated and the residue left to dry under vacuum overnight. The crude hydrochloride salt was used without further purification.
In a separate flask, alcohol **20** (57.9 mg, 105 μmol) and $Et_3N$ (41.0 μL, 294 μmol) were stirred at rt in anhydrous MeCN (10 mL) while solid DSC (75.3 mg, 294 μmol) was added. The mixture was stirred overnight, after which additional $Et_3N$ (22 μL, 157 μmol) was added. A solution of the amine in DMSO (1 mL) was added and the mixture stirred until consumption of starting material. The volatiles were removed under vacuum and the residue purified by reverse phase HPLC eluting with a gradient of MeOH in $H_2O$ (50 – 95%). The proximity probe **PP1** (7.60 mg, 6%) was isolated as a yellow solid and characterized as mixture of diastereomers. $^1$H NMR (500 MHz, $CD_3OD$) $\delta$ $^1$H NMR (500 MHz, $CD_3OD$) $\delta$ 7.78 (s, 2H), 7.58 (s, 2H), 7.44 (d, $J$ = 7.7 Hz, 4H), 7.29 – 7.23 (m, 4H), 7.17 (d, $J$ = 1.7 Hz, 1H), 6.25 (q, $J$ = 6.3 Hz, 1H), 5.51 (s, 4H), 4.51 – 4.41 (m, 3H), 4.37 (s, 4H), 4.31 – 4.21 (m, 3H), 4.06 (q, $J$ = 6.0 Hz, 4H), 4.00 – 3.88 (m, 8H), 3.23 – 2.96 (m, 19H), 2.94 – 2.84 (m, 3H), 2.73 – 2.63 (m, 3H), 2.46 (t, $J$ = 7.3 Hz, 4H), 2.22 – 2.09 (m, 8H), 2.10 – 1.98 (m, 4H), 1.78 – 1.16 (m, 35H), 0.98 (d, $J$ = 20.3 Hz, 7H). $^{13}$C NMR (125 MHz, $CD_3OD$) $\delta$ 175.97, 175.96, 175.90, 175.89, 175.14, 175.13, 174.61, 174.61, 174.44, 174.37, 170.32, 166.08, 166.07, 161.11, 160.78, 157.44, 157.42, 155.72, 155.62, 148.67, 148.58, 141.02, 140.73, 139.89, 139.89, 139.87, 137.22, 135.31, 135.15, 135.15, 129.63, 129.59, 128.60, 114.45, 110.16, 110.13, 109.45, 109.40, 70.23, 70.12, 69.62, 68.47, 63.35, 63.34, 61.60, 61.59, 57.00, 56.95, 56.39, 49.85, 49.51, 49.34, 49.17, 49.00, 48.83, 48.66, 48.49, 43.88, 41.05, 40.21, 40.18, 40.11, 40.09, 36.82, 36.80, 33.41, 33.02, 32.88, 31.54, 31.48, 31.36, 31.32, 29.96, 29.85, 29.77, 29.48, 26.91, 26.88, 26.41, 26.36, 25.13, 24.99, 24.31, 24.27, 22.45, 22.40, 19.74, 19.73. HRMS (ESI$^+$) calculated for $C_{53}H_{73}N_{15}O_{12}SH$ [M + H]$^+$ 1144.5362, found 1144.5371.

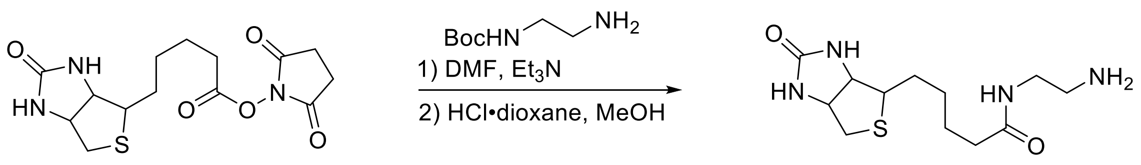


**21** **22**

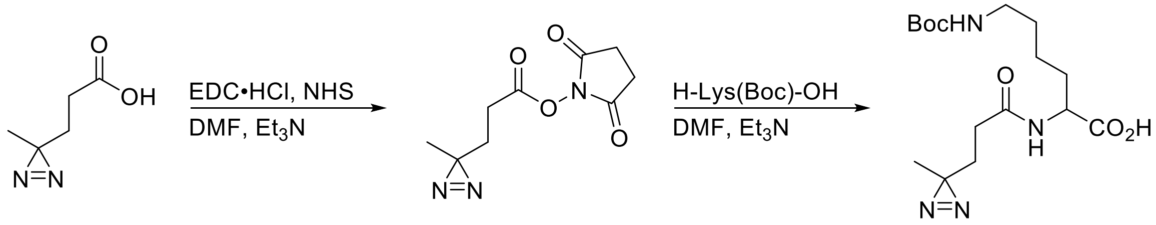


**14** **23** **24**

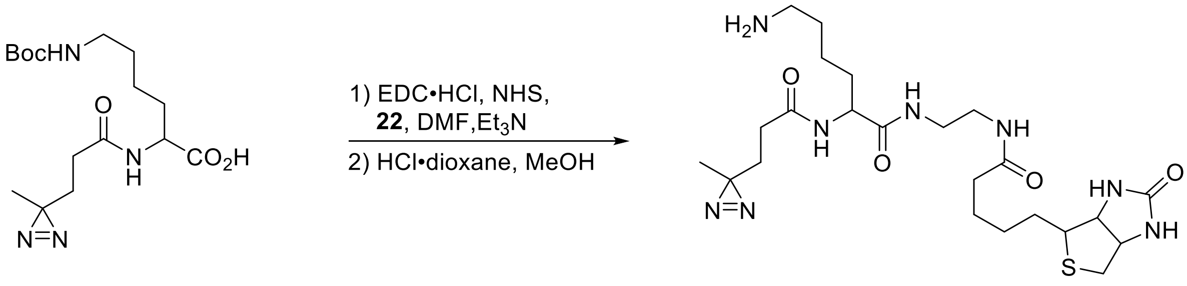


**24** **26**

**(2,5-dioxopyrrolidin-1-yl)5-(2-oxo-1,3,3a,4,6,6a-hexahydrothieno[3,4-d]imidazol-4-yl)pentanoate (21)**

An oven dried 100 mL round-bottom containing D-biotin (2.44 g, 10 mmol), EDC•HCl (2.87 g, 15 mmol), and NHS (1.73 g, 15 mmol) in 40 mL of anhydrous DMF was stirred overnight at rt. Upon consumption of starting material, cold $Et_2O$ was flowed in and supernatant decanted from the resultant oil. The residue was sonicated in fresh $Et_2O$ to provide pure NHS-ester **21** (5.73 g, 83%) as a fine white powder that was isolated by vacuum filtration and dried under suction. $^1H$ NMR (500 MHz, DMSO-$d^6$) $\delta$ 6.45 (s, 1H), 6.38 (s, 1H), 4.31 (dd, $J$ = 7.6, 5.0 Hz, 1H), 4.15 (ddd, $J$ = 7.7, 4.5, 1.8 Hz, 1H), 3.10 (ddd, $J$ = 8.3, 6.3, 4.3 Hz, 1H), 2.87 – 2.79 (m, 5H), 2.67 (t, $J$ = 7.4 Hz, 2H), 2.58 (d, $J$ = 12.4 Hz, 1H), 1.71 – 1.58 (m, 3H), 1.55 – 1.34 (m, 3H). $^{13}C$ NMR (125 MHz, DMSO-$d^6$) $\delta$ 170.35, 169.01, 162.80, 61.07, 59.24, 55.31, 30.06, 27.89, 27.65, 25.50, 24.37.

***N*-(2-aminoethyl)-5-(2-oxo-1,3,3a,4,6,6a-hexahydrothieno[3,4-d]imidazol-4-yl)pentanamide (22)**

Neat tert-butyl *N*-(2-aminoethyl)carbamate was added to a suspension of NHS-ester **21** in anhydrous DMF (8.0 mL) and the reaction stirred at rt overnight. Upon consumption of starting material $Et_2O$ was flowed in the mixture chilled to -20 °C overnight. The resultant precipitate was collected via vacuum filtration and dried under suction to yield Boc-protected amine (1.18 g, 78%) as a white powder. The intervening carbamate was stirred in MeOH containing 4M HCl to provide the amine after evaporation of the solvent, which was taken on without further purification. $^1H$ NMR (500 MHz, DMSO-$d^6$) $\delta$ 6.45 (s, 1H), 6.39 (s, 1H), 4.30 (dd, $J$ = 7.7, 5.1 Hz, 1H), 4.13 (ddd, $J$ = 7.8, 4.4, 1.9 Hz, 1H), 3.28 (q, $J$ = 6.2 Hz, 2H), 3.10 (ddd, $J$ = 8.6, 6.1, 4.4 Hz, 1H), 2.86 – 2.78 (m, 3H), 2.57 (d, $J$ = 12.4 Hz, 1H), 2.10 (t, $J$ = 7.5 Hz, 2H), 1.65 – 1.41 (m, 4H), 1.39 – 1.22 (m, 2H). $^{13}C$ NMR (125 MHz, DMSO-$d^6$) $\delta$ 172.78, 162.75, 61.05, 59.22, 55.42, 38.59, 36.39, 35.16, 28.26, 28.08, 25.03.

**(2,5-dioxopyrrolidin-1-yl) 3-(3-methyldiazirin-3-yl)propanoate (23)**
To a solution of crude acid **14** (2.40 g, 18.7 mmol) in anhydrous DMF (37 mL) was added EDC•HCl (4.48 g, 23.4 mmol) and NHS (2.69 g, 23.4 mmol). This mixture was stirred at room temperature overnight. Upon completion the reaction was diluted with $H_2O$ and extracted with $Et_2O$ (50 mL x 3). The combined extracts were dried over $Na_2SO_4$, filtered, and concentrated under vacuum. Precipitate that formed during evaporation of the volatiles was filtered and dried under suction to provide NHS ester **23** (2.27 g, 54%) as a white solid. $^1$H NMR (500 MHz, $CDCl_3$) *δ* 2.83 (s, 4H), 2.54 – 2.47 (m, 2H), 1.86 – 1.75 (m, 2H), 1.06 (s, 3H). $^{13}$C NMR (125 MHz, $CDCl_3$) *δ* 169.08, 167.71, 29.58, 25.82, 25.68, 24.86, 19.59.

**Biotinyl *N*-Boc-Photo Lysine (26)**
$N^6$-(tert-butoxycarbonyl)-L-lysine (246 mg 1.00 mmol) and $Et_3N$ (278 µL, 2.00 mmol) in anhydrous DMF (4 mL) were stirred while solid NHS-ester **23** (270 mg, 1.2 mmol) was added. The reaction was stirred at rt and progress monitored by LC/MS. Upon consumption of starting material, EDC•HCl (287 mg, 1.50 mmol) and NHS (173 mg, 1.50 mmol) were added at rt. The mixture was stirred at rt overnight, after which chilled $Et_2O$ was flowed in and resultant precipitate isolated by vacuum filtration. The NHS ester (319 mg, 0.704 mmol) and $Et_3N$ (293 µL, 2.11 mmol) were combined in fresh anhydrous DMF (4 mL) with stirring while amine **22** (242 mg, 0.845 mmol) was added. Upon completion, diethyl ether was flowed in and the reaction stored at -20 °C overnight. The resultant precipitate was isolated by vacuum filtration and dried under suction to afford carbamate **26** (323 mg, 73%) as a beige solid. $^1$H NMR (500 MHz, $CD_3OD$) *δ* 4.51 (dd, *J* = 7.9, 4.8 Hz, 1H), 4.38 – 4.30 (m, 1H), 4.25 – 4.17 (m, 1H), 3.32 – 3.18 (m, 2H), 3.09 – 2.98 (m, *J* = 6.8 Hz, 3H), 2.93 (dd, *J* = 12.8, 4.9 Hz, 1H), 2.72 (d, *J* = 12.7 Hz, 1H), 2.26 – 2.09 (m, 4H), 1.89 – 1.56 (m, 7H), 1.52 – 1.41 (m, 18H), 1.02 (d, *J* = 2.2 Hz, 3H). $^{13}$C NMR (125 MHz, $CD_3OD$) *δ* 176.29, 175.43, 174.83, 174.76, 174.63, 174.59, 174.51, 166.03, 158.42, 79.77, 63.22, 63.15, 61.61, 56.89, 55.05, 55.02, 53.50, 41.03, 40.17, 39.91, 39.85, 36.78, 36.73, 32.56, 32.20, 31.38, 31.22, 31.19, 30.99, 30.95, 29.68, 29.57, 29.42, 28.80, 28.78, 26.74, 26.38, 26.29, 24.21, 24.13, 19.77, 19.69.

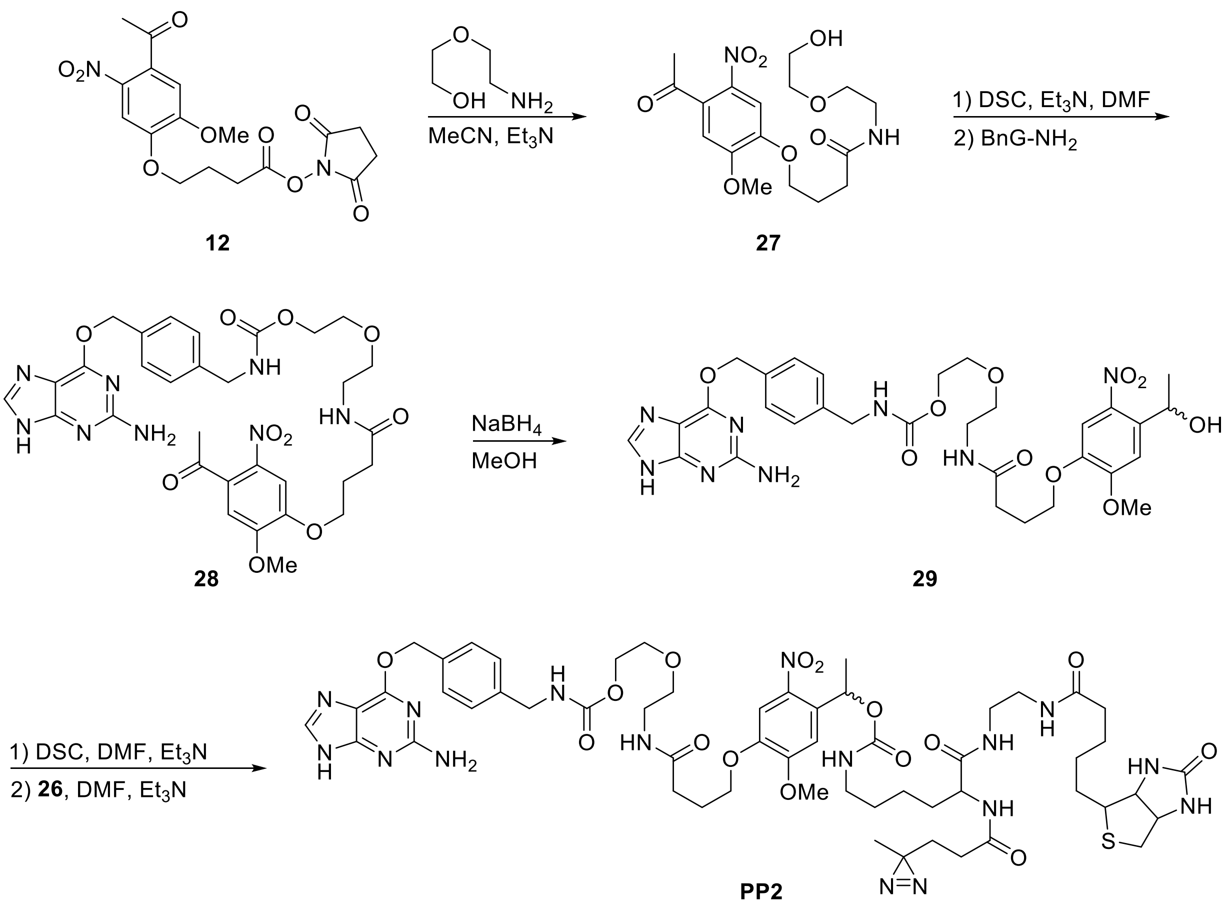


**4-(4-acetyl-2-methoxy-5-nitrophenoxy)-*N*-[2-(2-hydroxyethoxy)ethyl]-butanamide**
Neat 2-(2-aminoethoxy)ethanol (158 µL, 1.50 mmol) was added to a suspension of NHS-ester **12** (394 mg, 1.00 mmol) and $Et_3N$ (150 µL, 1.00 mmol) in anhydrous MeCN (5 mL). The reaction was stirred at rt for 5 hours. The volatiles were removed under vacuo and the residue purified by reverse-phase HPLC, eluting with a gradient of MeOH in $H_2O$ (40–95%). Pure amide **27** (248 mg, 64%) was isolated as a yellow oil. $^1H$ NMR (500 MHz, $CDCl_3$) $\delta$ 7.57 (s, 1H), 6.73 (s, 1H), 4.11 (t, $J$ = 6.1 Hz, 2H), 3.93 (s, 3H), 3.71 (t, $J$ = 4.4 Hz, 2H), 3.53 (t, $J$ = 5.1 Hz, 4H), 3.45 (q, $J$ = 5.1 Hz, 2H), 2.46 (s, 3H), 2.44 (t, $J$ = 7.3 Hz, 2H), 2.18 (p, $J$ = 6.7 Hz, 2H). $^{13}C$ NMR (125 MHz, $CDCl_3$) $\delta$ 200.48, 154.27, 148.83, 138.35, 132.84, 108.79, 108.16, 77.36, 72.15, 69.69, 68.63, 56.69, 39.63, 32.56, 30.46, 24.93.

**Alcohol (29)**
In an oven dried flask, DSC (316 mg, 1.23 mmol) was added to an anhydrous DMF (3 mL) solution of amide **27** (434 mg, 1.13 mmol) and $Et_3N$ (428 µL, 3.08 mmol). The reaction was stirred overnight at room temperature before adding solid amine **7** (278 mg, 1.03 mmol). Stirring was continued at rt for an additional 3 hours. Upon completion, the reaction was diluted with $H_2O$ and extracted with EtOAc. The combined organics were washed with brine, dried over $Na_2SO_4$, filtered, and solvent removed under vacuum to provide the methyl ketone (467 mg, 61%). $^1H$ NMR (500 MHz, $CD_3OD$) $\delta$ 7.80 (s, 1H), 7.45 (s, 1H), 7.38 (d, $J$ = 7.8 Hz, 2H), 7.31 (s, 1H), 7.22 (d, $J$ = 7.8 Hz, 2H), 5.41 (s, 2H), 4.24 (s, 2H), 4.16 – 4.10 (m, 2H), 3.95 (t, $J$ = 6.3 Hz, 1H), 3.87 (s, 2H), 3.59 (t, $J$ = 4.8 Hz, 2H), 3.49 (t, $J$ = 5.4 Hz, 2H), 2.34 (t, $J$ = 7.5 Hz, 2H), 2.03 (h, $J$ = 6.6 Hz, 2H), 1.43 (d, $J$ = 6.3 Hz, 3H). $^{13}C$ NMR (125 MHz, $CD_3OD$) $\delta$ 175.39, 161.50,

158.97, 155.23, 148.02, 140.48, 138.93, 136.65, 129.54, 128.25, 109.79, 70.39, 70.35, 69.49, 68.64, 66.21, 65.02, 56.66, 45.14, 40.36, 36.94, 33.36, 31.63, 26.35, 25.16.
Subsequently, the ketone was taken up in MeOH (10 mL) and solid $NaBH_4$ added in portions (77.9 mg, 2.06 mmol) and vigorously stirred at rt. Once the intervening ketone was consumed the reaction was quenched with 10 mL $H_2O$ and stirred for an additional 30 min. The mixture was then extracted with EtOAc, organics dried over $Na_2CO_3$, filtered, and concentrated to provide Alcohol **29** that was taken on without further purification.

**Photoproximity probe 2 (PP2)**
A solution of carbamate **26** (115 mg, 205 µmol) in MeOH (1 mL) was added to 4 M HCl•dioxane (2 mL) at 0 °C. The temperature was allowed to come to rt while stirring for 2 hrs. Upon disappearance of starting material the volatiles were evaporated and the residue left to dry under vacuum overnight. The crude hydrochloride salt was used without further purification.
In a separate flask, alcohol **29** (70.1 mg, 102 µmol) and *i*$Pr_2NEt$ (68.0 µL, 411 µmol) were stirred at rt in anhydrous DMF (1 mL) while solid DSC (52.5 mg, 205 µmol) was added. The mixture was stirred overnight. A solution of the amine in DMSO (1 mL) was added and the mixture stirred until consumption of starting material. The volatiles were removed under vacuum and the residue purified by reverse phase HPLC eluting with a gradient of MeOH in $H_2O$ (50 – 95%). The proximity probe **PP2** (12.8 mg, 10%) was isolated as a yellow solid and characterized as mixture of diastereomers. $^1H$ NMR (500 MHz, DMSO-$d^6$) $\delta$ 8.04 (s,1H), 7.96 (t, J = 5.6 Hz, 2H), 7.80 (s, 1H), 7.56 (s, 1H), 7.45 (d, *J* = 7.9 Hz, 2H), 7.27 (d, *J* = 7.7 Hz, 2H), 7.11 (s, 1H), 6.46 (d, *J* = 9.1 Hz, 1H), 6.38 (s, 1H), 6.21 (s, 2H), 6.10 (q, *J* = 6.5 Hz, 1H), 5.45 (s, 2H), 4.35 – 4.28 (m, 1H), 4.18 (d, *J* = 6.3 Hz, 2H), 4.15 – 4.02 (m, 5H), 3.90 (s, 3H), 3.58 – 3.54 (m, 2H), 3.44 – 3.40 (m, 2H), 3.26 – 3.17 (m, 5H), 3.09 (d, *J* = 15.4 Hz, 8H), 2.99 – 2.79 (m, 3H), 2.58 (d, *J* = 12.5 Hz, 2H), 2.25 (t, *J* = 7.5 Hz, 3H), 2.09 – 2.01 (m, 7H), 1.95 (q, *J* = 6.9 Hz, 2H), 1.67 – 1.41 (m, 7H), 1.41 – 1.15 (m, 3H), 1.00 – 0.96 (m, 4H). $^{13}C$ NMR (125 MHz, DMSO-$d^6$) $\delta$ 178.40, 178.33, 172.32, 171.87, 171.57, 170.97, 170.86, 162.76, 161.14, 160.91, 159.44, 156.44, 155.13, 153.56, 153.40, 146.76, 139.62, 139.24, 135.34, 133.54, 133.42, 128.52, 127.81, 127.08, 108.39, 69.05, 68.65, 68.32, 67.02, 66.51, 63.23, 61.03, 59.22, 56.19, 55.42, 55.35, 52.59, 43.57, 38.49, 38.18, 35.23, 31.61, 31.50, 29.78, 29.59, 29.08, 28.22, 28.06, 25.85, 25.20, 24.64, 22.70, 21.94, 19.29. HRMS (ESI$^+$) calculated for $C_{55}H_{76}N_{16}O_{15}SH$ $[M + H]^+$ 1233.5475, found 1233.5480.

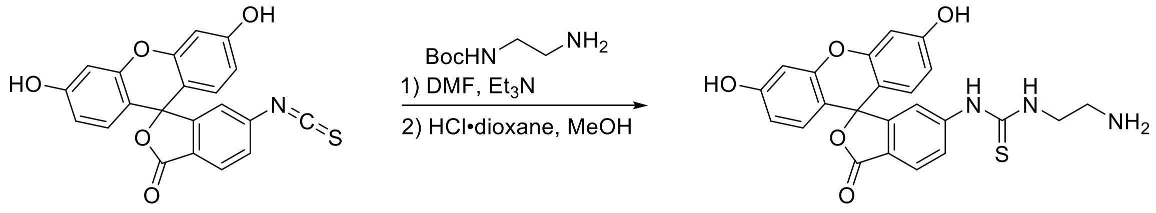


**30** **31**

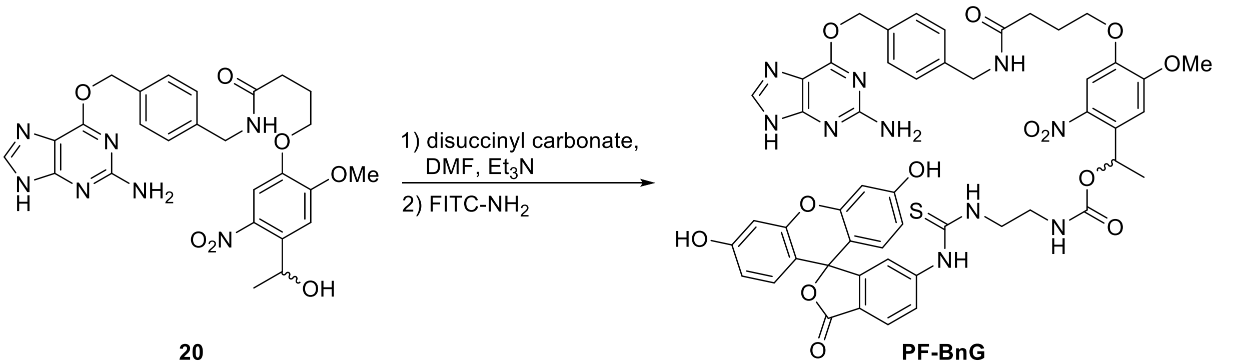


**20** **PF-BnG**

**Photocleavable FITC-benzyl guanine (PF-BnG)**
Fluorescein isothiocyanate (**FITC**, 100 mg, 0.257 mmol) was added to a DMF solution (1 mL) of *N*-Boc-ethylenediamine (41.3 µL, 0.262 mmol) and $Et_3N$ (7.2 µL, 51.4 µmol) then stirred at room temperature overnight. Upon completion the volatiles were remove under vacuum and the residue purified by reverse-phase HPLC eluting with a gradient of MeOH in $H_2O$. The purified carbamate(102 mg, 72%) was isolated as an orange solid and used directly.
The intervening carbamate (73.4 mg, 133 µmol) was stirred for 1 hour at room temperature in MeOH (2 mL), 4M HCl•dioxane (0.5 mL), and triisopropylsilane (47 µL). Once fully deprotected the volatiles were removed under vacuum and the residue used without further purification.
In a separate flask, *N,N'*-disuccinimidyl carbonate (42 mg, 163 µmol) was added to a DMF (1 mL) solution of Alcohol **20** (60.4 mg, 109 µmol) and *i*$Pr_2NEt$ (96 µL, 582 µmol). The mixture was left stir at room temperature overnight. The mixture was transferred to the above amine hydrochloride and stirred at room temperature. Upon completion, the solvent was removed, and the residue purified by reverse-phase HPLC eluting with a gradient of MeOH in $H_2O$ to provide **PF-BnG** (36 mg, 22%) as an orange-solid. $^1H$ NMR (500 MHz, $CD_3OD$) $\delta$ 8.03 (s, 1H), 7.95 (s, 1H), 7.67 (dd, *J* = 8.4, 2.0 Hz, 1H), 7.49 (s, 1H), 7.41 (d, *J* = 8.1 Hz, 2H), 7.25 (d, *J* = 8.0 Hz, 2H), 7.14 (s, 1H), 7.08 (d, *J* = 8.3 Hz, 1H), 6.73 – 6.62 (m, 4H), 6.53 (ddd, *J* = 8.7, 4.0, 2.4 Hz, 2H), 6.28 (q, *J* = 6.3 Hz, 1H), 5.53 (s, 2H), 4.35 (s, 2H), 4.03 – 3.91 (m, 2H), 3.89 (s, 3H), 3.64 – 3.55 (m, 1H), 3.45 – 3.33 (m, 1H), 3.30 – 3.24 (m, 2H), 2.41 (t, *J* = 7.3 Hz, 2H), 2.07 (p, *J* = 6.8 Hz, 2H), 1.55 (d, *J* = 6.4 Hz, 3H). $^{13}C$ NMR (125 MHz, $CD_3OD$) $\delta$ 183.20, 175.18, 171.08, 161.38, 161.23, 160.21, 158.32, 155.63, 155.53, 154.13, 148.57, 140.85, 140.38, 136.20, 135.22, 130.42, 130.30, 129.88, 128.92, 128.66, 113.60, 111.43, 110.11, 109.38, 103.51, 69.87, 69.56, 69.50, 57.01, 43.83, 33.38, 26.34, 22.54. HRMS ($ESI^+$) calculated for $C_{50}H_{46}N_{10}O_{13}SH$ $[M + H]^+$ 1027.3044, found 1027.3024.

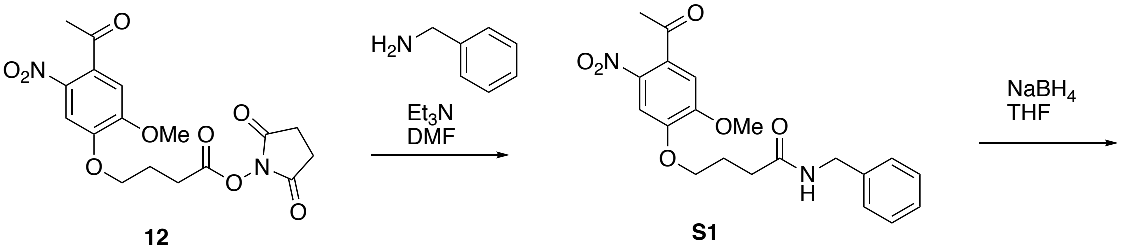


**12** **S1**

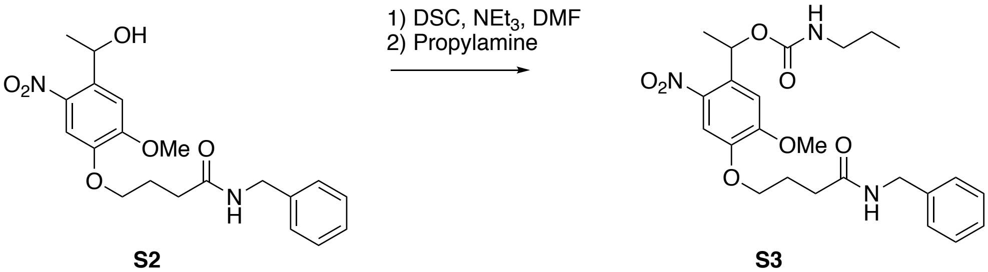


**S2** **S3**

**Ketone (S1).** Benzylamine (.83 mL, 7.6 mmol) was added dropwise to a solution of NHS-ester **12** (1.00 g, 2.53 mmol) in DMF (25 mL) under nitrogen. TEA (.4 mL, 2.8 mmol) was added dropwise and the reaction was allowed to stir overnight. Upon completion, the reaction was chilled in an ice bath and deionized water was added. After sonication, the resulting precipitate was vacuum filtered and washed with deionized water and cold $Et_2O$ to yield pure ketone **S1** (.92 g, 94%) as a beige powder. 1H NMR (500 MHz, $CD_3O_D$) δ 7.59 (s, 1H), 7.27-7.28 (m, 5H), 7.16 (s, 1H), 6.28-6.31 (q, 1H), 4.38 (s, 2H), 4.09 (t, 2H), 3.93 (s, 3H), 2.67 (s, 3H), 2.48 (t, 2H), 2.13 (m, 2H).

**Alcohol (S2).** Solid $NaBH_4$ (125 mg, 3.30 mmol) was added in portions to a solution of S1 (440 mg, 1.14 mmol) in THF (23 mL) at room temperature and was monitored via TLC. After stirring for 24 hours, the reaction was brought to 40 °C and allowed to stir overnight. Upon completion, the reaction was quenched with $NH_4Cl$ and extracted with ethyl acetate (100 mL x 3). The combined organic layers were washed with NaCl and the solution was dried with $Na_2SO_4$. Rotary evaporation yielded crude alcohol as a yellow oil which was then purified via column chromatography on silica gel (DCM:MeOH;100:1 to 20:1) to yield pure alcohol as a light-yellow solid **S2** (.41 g, 93%) 1H NMR (500 MHz, $CDCl_3$) δ 7.59 (s, 1H), 7.27-7.28 (m, 5H), 7.16 (s, 1H), 6.28-6.31 (q, 1H), 4.46-4.48 (d, 2H), 4.18 (t, 2H), 3.91 (s, 3H), 2.52 (s, 3H), 2.49 (t, 2H), 2.27 (m, 2H).

**Nitroveratryl Carbamate Model Linker** (**S3**). A DMF (5.4 mL) solution of alcohol **S2** (104 mg, .27 mmol) was prepared and DSC (138 mg, .54 mmol) was added at room temperature under nitrogen and was allowed to stir for 15 h. Propylamine (.13 mL, 1.62 mmol) was then added dropwise and the reaction was brought to 34 °C and allowed to stir for 24 hr. Upon completion, the reaction was quenched with cold deionized water and extracted with ethyl acetate (100 mL x 3). The combined organic layers were washed with NaCl and the solution was dried with $Na_2SO_4$. Rotary evaporation yielded crude nitroveratryl carbamate as a yellow oil which was then purified via column chromatography on silica gel (DCM:MeOH; 200:1 to 20:1) to yield pure nitroveratryl carbamate **S3** as a white solid (52 mg, 41%). 1H NMR (500 MHz, $CD_3OD$) δ 7.60

(s, 1H), 7.27 (m, 5H), 7.18 (s, 1H), 6.26-6.30 (q, 1H), 4.38 (s, 2H), 4.09 (t, 2H), 3.93 (s, 3H), 3.02 (s, 3H), 2.48 (t, 2H), 2.16 (m, 2H), 1.60-1.61(d, 3H), 1.49 (m, 2H), .89 (t, 3H). 13C NMR (125 MHz, $CD_3OD$) $\delta$ 171.85, 155.62, 154.02, 147.22, 140.09, 139.73, 133.99, 128.70, 127.63, 127.16, 108. 91, 108.81, 68.74, 67.46, 56.63, 42.50, 42.33, 32.00, 25.15, 23.06, 22.32, 11.61.

**General biological methods**

**Cell culture**

HEK293T cell lines were purchased from ATCC and all HEK293T lines were propagated in DMEM (Corning) supplemented with 10% fetal bovine serum (FBS, Corning) and 1% penicillin/streptomycin (Gibco). All cell lines were grown at 37 ˚C in a 5% $CO_2$ humidified incubator.

**SDS-PAGE and Western blot**

Cells were harvested by scraping, pelleted by centrifugation, washed twice with PBS and lysed in 8 M urea, 50 mM $NH_4HCO_3$ and EDTA-free complete protease inhibitor (Roche), pH 8.0, at 4 ˚C. Cells were sonicated (Fisher Scientific FB-505), insoluble debris cleared by centrifugation, and the supernatant was diluted into 4X Laemmli buffer containing 50 mM dithiothreitol (DTT) or 6% beta-mercaptoethanol (βME) as reducing agents. Samples were prepared for SDS-PAGE by heating to 95 ˚C for 5 minutes, cooled to room temperature, resolved on NuPAGE Novex 4-12% Bis-Tris Protein Gels (Invitrogen) or 10% SDS-PAGE gel, and transferred to nitrocellulose membranes by standard western blotting methods. Membranes were blocked in 2% BSA in TBS containing 0.1% tween-20 (TBST) and probed with primary and secondary antibodies. Primary antibodies used in this study include: anti-FLAG-M2 (1:1000, F1804, Sigma Aldrich), Streptavidin-IRdye800 (1:10,000, 92632230, Li-cor). Secondary donkey anti-rabbit, donkey anti-goat, and donkey anti-mouse (Licor), were used at 1:10,000 dilution in 2% BSA-containing TBST and incubated for 1 hour prior to washing and imaging on a Licor infrared scanner. Densitometry measurements were performed with ImageJ software.

**Circular polymerase extension cloning (CPEC) construction of mammalian plasmids**

All PCR reactions were performed using NEB Q5 high-fidelity polymerase (M0491S) and Promega dNTP mix (U1515). Initial constructs were generated using the pSnapf vector from NEB (#N9183S), the pFlag-Keap1 vector from Addgene (#28023) along with the following CPEC primers purchased from IDT:

*pDC-002 (Keap1-SNAP-Flag3x):*
F1 = ATGGACAAAGACTGCGAAATGAAGCGCACCACCC
R1 = ACCCAGCCCAGGCTTGCCCA
F2 = GGCAAGCCTGGGCTGGGTgactacaaagaccatgacggtgattataaagatcatgacat
R2 = GTGCGCTTCATTTCGCAGTCTTTGTCCATGCTTCCGCCGCCgcggccgccacaggtaca
*pDC-003 (SNAP-Keap1-Flag3x):*
F1 = atgcagccagatcccaggcctagc
R1 = gatatctgcagaattccaccacactggactagtggatcc
F2 = ccagtgtggtggaattctgcagatatcATGGACAAAGACTGCGAAATGAAGCGCACCAC
R2 = gcctgggatctggctgcatGCTCCCTCCGCCGCCACCCAGCCCAGGCTTGCCC

The constructs above were subcloned into the pLenti6/V5-p53_R273H Addgene (#22934) vector for lentiviral transduction using the following CPEC primers purchased from IDT:

*pDC-006 (Keap1-SNAP-Flag3x):*
F1 = tagtaatgagtttggaattaattctgtggaatgtgtgtcagttaggg

R1 = ggtgaagggatcaattccaccacactgg
F2 = GGTGGAATTGATCCCTTCACCatgcagccagatcccagg
R2 = cacattccacagaattaattccaaactcattactacttgtcatcgtcatccttgtagtcg
*pDC-007 (SNAP-Keap1-Flag3x):*
F1 = tagtaatgagtttggaattaattctgtggaatgtgtgtcagttaggg
R1 = ggtgaagggatcaattccaccacactgg
F2 = ggtggaattgatcccttcaccATGGACAAAGACTGCGAAATGAAGC
R2 = cacattccacagaattaattccaaactcattactacttgtcatcgtcatccttgtagtcg

**Transient and Stable Protein Expression in Cells**
Mammalian cells stably expressing the KEAP1 SNAP-Tag fusions were obtained by co-transforming a 6 cm plate of HEK293T cells with 0.1 µg pCMV-VSV-G (Addgene #8454),0.9 µg pCMV delta R8.2 (Addgene #12263), and 1.0 µg of either pDC-006 or pDC-007. The resultant viral media was collected at 24 and 48 hours, passed through a 0.45-micron filter, and diluted with serum-free DMEM containing 8 µg/mL polybrene (Sigma), final concentration. Viral transduction was achieved by culturing a separate population of HEK293T in the diluted viral media for 24 hours. Afterward, the viral media was removed and the transduced cells grown in full DMEM containing 8 µg/mL blasticidin (Gibco). Stable incorporation of the transgene was confirmed by western blotting using monoclonal anti-FLAG M2 antibody (Sigma). Stable expression of Snap-Flag, derived from pSnapf vector (NEB #N9183S), was achieved through chemical selection of transiently transfect HEK293T cells with 500 µM G418 (Gibco). Transient SNAP-FLAG protein expression was accomplished using transfection with lipofectamine 2000 according to manufacturer protocol.

**In vitro photocleavage assay**
Recombinant purified SNAP-Tag (20 µg) was incubated in 200 µL DPBS containing 5 µM **PF-BnG** for 1 hour at 37 °C. Afterward, the reaction was divided into six 30 µL aliquots, placed on ice, and irradiated with 365 nm light using a spectrolinker XL-1500a (Spectroline). Samples were successively removed from irradiation after 0, 0.5, 1, 2.5, 5, 10 minutes of irradiation, diluted with loading buffer, and run on SDS-PAGE gel. Photocleavage was judged by in-gel fluorescence visualized using a ChemiDoc imaging system (Bio-Rad) scanning for fluorescein.

**SNAP-flag-NLS plasmid generation**

Consecutive rounds of site-directed mutagenesis PCR reactions were performed with the Tag-Master Kit (GM Bioscience, #7002) using the pSnapf vector from NEB (#N9183S) and the following primers purchased from IDT:
SNAP-flag_for
ggcggatccgcgtttGATTACAAGGATGACGACGATaaactcgaggttaattaa
SNAP-flag_rev
ttaattaacctcgagtttATCGTCGTCATCCTTGTAATCaaacgcggatccgcc
SNAP-flag-NLS_for
gataaactcgaggttaatCCGAAGAAGAAGCGCAAGGTGtaatgagcggccgca
SNAP-flag-NLS_rev
tgcggccgctcattaCACCTTGCGCTTCTTCTTCGGattaacctcgagtttatc

**Nitroveratryl Carbamate Linker and Diazirine Photochemical Kinetic LC-MS Analysis.** A .2 mM solution of nitroveratryl carbamate **S3** in methanol was prepared from a 10 mM DMSO stock solution. 1 mL of the solution was added to three individual Eppendorf tubes for three replicates. The samples were placed on ice and positioned approximately 4 cm the source of irradiation. The samples were irradiated with 365 nm light using a spectrolinker XL-1500a (Spectroline) for several time points and analyzed via LC-MS on an Agilent 1200 Series G1311A using the following method [Buffer A: 95% $H_2O$, 5% MeCN, .1% TFA. Buffer B: 95% MeCN, 5% $H_2O$, .1% TFA. 0 min (85% A, 15% B), 2min (85% A, 15% B), 2.5 min (60% A, 40% B), 8 min (0% A, 100% B), 9.75 min (0% A, 100% B), 10 min (85% A, 15% B), 12 min (85% A, 15% B) at .5 mL/min. Integration data for UV (215 nm) were used to derive kinetic parameters on Graph Pad Prism 8. For the diazirine model compound, .2 mM solution of diazirine model **15** in methanol was prepared from a 10 mM DMSO stock solution. 1 mL of the solution was added to three individual Eppendorf tubes for three replicates. The samples were placed on ice and positioned approximately 4 cm the source of irradiation. The samples were irradiated with 365 nm light using a spectrolinker XL-1500a (Spectroline) for several time points and analyzed via LC-MS 1200 Series G1311A using the following method [Buffer A: 95% $H_2O$, 5% MeCN, .1% TFA. Buffer B: 95% MeCN, 5% $H_2O$, .1% TFA. Gradient: 0 min (75% A, 25% B), 2min (75% A, 25% B), 2.5 min (60% A, 40% B), 8 min (0% A, 100% B), 9.75 min (0% A, 100% B), 10 min (75% A, 25% B), 12 min (75% A, 25% B) at .5 mL/min. Integration data for UV (215 nm) was used to calculate kinetic parameters on Graph Pad Prism 8.

**Fluorescence microscopy validation of SNAP-flag-NLS nuclear-localization**

HeLa cells were transiently-transfected with the SNAP-flag-NLS plasmid using Lipofectamine 2000 (ThermoFisher Scientific, 11668019) following the manufacturer's protocol. Following overnight recovery, transfected cells were trypsinized and seeded at 20,000 cells per well in a 12-well chamber slide (Ibidi, #81201). Following overnight incubation, the media was removed, cells were washed with PBS three times, fixed using 4% PFA/PBS for 10 min, washed with PBS three time, permeabilized with 1% Triton-X-100/PBS for 1 min, blocked with 2% BSA/PBS-T for 1 hr, primary-stained overnight with a 1:200 dilution of anti-FLAG antibody in 2% BSA/PBS-T at 4°C with gentle rocking, washed 3x 5 min with PBS-T, secondary-stained with a 1:500 dilution of anti-mouse-Alexa488 (ThermoFisher Scientific, #A-11001) in 2% BSA/PBS-T for 1 hr, washed 3x 5 min with PBS-T, counter-stained with 1:5000 DAPI in PBS for 20 min, washed 2x 5 min with PBS, and then slides were sealed. Fluorescent images were captured with an Olympus DSU spinning disk confocal microscope (Olympus Corporation of the Americas, Center Valley, PA) with a Hamamatsu model C9100 EM-CCD camera (Hamamatsu Photonics, Skokie, IL) run by SlideBook v5.0 software (Intelligent Imaging Innovations, Denver, CO).

**Fluorescence microscopy of PF-BnG nuclear localization and photo-cleavage**

HeLa cells were transiently-transfected with the SNAP-flag-NLS plasmid using Lipofectamine 2000 (ThermoFisher Scientific, 11668019) following the manufacturer protocol. Following overnight recovery, transfected cells were trypsinized and seeded at 20,000 cells per well in a 12-well chamber slide (Ibidi, #81201). Following overnight incubation, media was removed and replaced with serum-free RPMI containing DMSO vehicle, 10 µM BnG-FITC, or 15 µM PF-BnG. Following a 2 hr incubation and subsequent 30 min probe soak out in full media, the plate was covered, put on ice, and wells were exposed to 0, 1, 5, or 10 min of UV irradiation. Following photo-cleavage, cells were washed with PBS three times, fixed using 4% PFA/PBS for 10 min, washed with PBS three times, and then slides were sealed. Fluorescent images were captured with an Olympus DSU spinning disk confocal microscope (Olympus Corporation of the Americas, Center Valley, PA) with a Hamamatsu model C9100 EM-CCD camera (Hamamatsu Photonics, Skokie, IL) run by SlideBook v5.0 software (Intelligent Imaging Innovations, Denver, CO).

**PP1 dose response and viability assay**

*Dose response* – Each well of a 6-well plate was seeded with 300,000 HEK293T cells stably expressing either SNAP-FLAG or KEAP-SNAP. After reaching ~90% confluency the growth media was removed, cells washed with DPBS, and treated with varying concentrations (0, 0.5, 1, 5, 15, 45 µM) of photoproximity probe **PP1** in 500 µL serum-free DMEM for 2 hours at 37 °C. Post treatment, media was aspirated and non-reacted probe washed out with 3 mL full DMEM over 40 minutes, changing the media twice. Cells were harvested, washed, and lysed in DPBS containing protease inhibitor using a tip sonicator. Lysate was normalized to 1 mg/mL then diluted with loading buffer and run on SDS-PAGE gel that was transferred by western blot to nitrocellulose. The nitrocellulose membrane was stained for biotin and Flag-Tag using streptavidin IR dye (Li-cor #926-32230) and anti-Flag (Sigma # F3165, Li-cor #926-68072). Fluorescence images were collected using an Odyssey infrared imager (Li-cor).

*Viability assay* –HEK293T cells stably expressing SNAP-FLAG were seeded with at 5,000 cells per well in 50 µL DMEM in a 96-well plate. 24 hours later, cells were treated with a varying concentration (0, 0.5, 1, 5, 15, 45 µM in sextuplicate) of photoproximity probe **PP1** in a total volume of 100 µL DMEM. After 2 hours at 37 °C, a 100 µL of CellTiter-Glo (Promega) was added and the plate imaged using a Synergy Neo HST plate reader (BioTek).

**Co-immunoprecipitation experiments.** FLAG-SNAP and FLAG-KEAP1 expressing cell cultures (10 cm plate) were lysed in PBS containing 1 mM DTT and protease inhibitors (Roche Complete, EDTA-free) at 4 °C by tip sonication. Lysates were cleared by centrifugation at 13,000 rpm, and immunoprecipitations were initiated by loading onto anti-FLAG resin (50 µL slurry) prewashed with lysis buffer, and incubated by rotating overnight at 4 °C. Beads were washed 4x 5 min with 500 µL buffer, and complexes eluted with 3xFLAG peptide (100 µg/mL) into SDS-loading buffer containing DTT. Input and enriched proteomes in SDS-loading buffer were boiled for 5 min at 95 °C and analyzed by SDS-PAGE gel and Western blot as indicated above. Western blots were stained for FLAG protein using anti-FLAG-M2 (1:1000, F1804, Sigma Aldrich), hexokinase 2 using anti-HK2 (1:1000, #2867, Cell Signaling) and secondary antibodies as indicated above.

**In vitro Proximity-labeling assays**

Clarified cell lysate from HEK293T cells either transiently (Fig 2b) or stably (Fig 2c, Fig. S4) expressing SNAP-Tag (NEB #N9183S) or SnapFlag were normalized to 1 mg/mL and 250 µL aliquots incubated in a microcentrifuge tube with either 500 nM of **PP1**, **PP2**, or DMSO for 1 hour at 37 °C. Washed anti-FLAG M2 affinity gel, 40 µL of a 50% slurry, was transferred to each reaction in 750 µL of DPBS and the resultant suspension left to rotate overnight at 4 °C (Sigma #A2220). Samples were then spun down at 4000 x g, supernatant aspirated, and the resin washed with 1M urea in DPBS (1 mL x 8) followed by DPBS (1 mL). After being resuspended in 100 µL of either DPBS the samples were placed on ice where they were irradiated with 365 nm light for 10 minutes (Spectrolinker XL-1500a). Once irradiated, the beads were washed with 100 mM pH 3.5 glycine buffer (100 µL x 2) and DPBS (1 mL). To elute the remaining SNAP-FLAG and resin-bound FLAG-antibody the beads were boiled for 5 minutes at 95 °C in 20 µL 4x-loading buffer containing 8% SDS and 400 mM DTT. After which, 60 µL of DPBS was added and the suspension boiled at 95 °C for an additional 5 minutes. The samples were then run on SDS-PAGE gel and transferred to nitrocellulose where they were stained with anti-mouse IR dye (Li-cor #926-68072) and streptavidin IR dye (Li-cor #926-32230). Bands corresponding to biotinylated proteins and the FLAG antibody were visualized using an Odyssey infrared imager (Li-cor).

**SILAC cell culture methods and proteomic sample preparation**

SILAC labeling was performed by growing cells for at least five passages in lysine- and arginine-free SILAC medium (RPMI, Invitrogen) supplemented with 10% dialyzed fetal calf serum, 2 mM L-glutamine and 1% Pen/Strep. “Light” and “heavy” media were supplemented with natural lysine and arginine (0.1 mg/mL), and $^{13}C$-, $^{15}N$-labeled lysine and arginine (0.1 mg/mL), respectively. General protein digestion for LC-MS/MS analysis was performed by diluting protein (e.g. whole lysate or enriched proteins) in digestion buffer (8 M urea, 50 mM $NH_4HCO_3$, pH 8.0), followed by disulfide reduction with DTT (10 mM, 40 minutes, 50 ˚C), alkylation (iodoacetamide, 15 mM, 30 min, room temperature, protected from light) and quenching (DTT, 5mM, 10 minutes, room temperature). The proteome solution was diluted 4-fold with ammonium bicarbonate solution (50 mM, pH 8.0), $CaCl_2$ added (1 mM) and digested with sequencing grade trypsin (~1:100 enzyme/protein ratio; Promega) at 37 ˚C while rotating overnight. Peptide digestion reactions were stopped by acidification to pH 2-3 with 1% formic acid, and peptides were then desalted on ZipTip C18 tips (100 µL, Millipore), dried under vacuum, resuspended with LC-MS grade water (Sigma Aldrich), and then lyophilized. Lyophilized peptides were dissolved in LC-MS/MS Buffer A ($H_2O$ with 0.1% formic acid, LC-MS grade, Sigma Aldrich) for proteomic analysis.

**Sample preparation and streptavidin enrichment of basal KEAP1 PhotoPPI profile.**

Quantitative proximity labeling study with SILAC quantitative proteomics was performed with “heavy” and “light” labeled HEK293T cells expressing KEAP SNAP fusion constructs. SILAC-labeled cells, grown to 80-90% confluency in 10 cm cell-culture treated plates (Denville) each, were incubated with DMSO alone (light cells) or PP1 probe (15 µM, heavy cells) for 2 hours in serum-free SILAC RPMI. After incubation, excess probe was washed out by media replacement and incubation for 10 min, performed twice. All cells were then incubated in 2 mL SILAC-RPMI, UV irradiated using a Spectroline XL-1500A instrument for 10 minutes, scraped, washed with cold PBS (2x), and tiny aliquots of cells (20 µL out of 500 µL resuspended cells in PBS) from each cell plate were taken for the analysis of bulk proteome. These “heavy” and “light” cell aliquots were combined and digested by general protein digestion protocol described above.

The rest of cells (480 µL out of 500 µL resuspended cells in PBS) were pelleted and then lysed in RIPA lysis buffer (50 mM Tris, 150 mM NaCl, 1% Triton X-100, 0.5% deoxycholate, pH 7.4) supplemented with EDTA-free complete protease inhibitor (Roche) and 1 mM DTT, at 4 ˚C. After sonication, insoluble debris was cleared by centrifugation (17,000 *g*, 10 min). Streptavidin C1 magnetic beads (30 µL slurry, 65001, Invitrogen) were washed twice with RIPA buffer, and each cell lysate was separately incubated with the magnetic beads with rotation overnight at 4 ˚C. The beads were subsequently washed five times with 0.5 mL of RIPA lysis buffer containing 1 mM DTT, combined together, then washed once with 1 mL of 1 M KCl, four times with 0.5 mL PBS, and two times with 2 M Urea in 25 mM ammonium bicarbonate. 500 µL of 6 M Urea in 50 mM ammonium bicarbonate was then added to the beads, and samples were reduced on resin by TCEP (10 mM final), with orbital shaking, for 20 minutes at 65 ˚C. Samples were then alkylated by adding iodoacetamide (20 mM final), covered from the light and with orbital shaking, for 40 minutes at 37 ˚C. The streptavidin magnetic beads were collected, washed once with 2 M Urea in 25 mM ammonium bicarbonate, and the buffer exchanged to 2 M Urea in 25 mM ammonium bicarbonate supplemented with 1 mM $CaCl_2$. Enriched proteins were digested on bead by the incubation of 2 µg sequencing grade trypsin overnight at 37 ˚C. Following trypsinization, supernatant was collected, acidified with HPLC grade formic acid (2% final, pH 2-3), and peptides were desalted as indicated above.

**Sample preparation for CBR470-1 perturbed KEAP1 PhotoPPI-profile**

HEK293T cells stably expressing Keap1 SnapTag fusions were cultured in 10 cm plates in SILAC RPMI. Once cultures reached ~60% confluency media was aspirated and both heavy and light conditions were treated with 10 µM CBR470 in 4 mL full SILAC RPMI. After a 14-hour incubation at 37 °C media was aspirated and replaced with 3 mL of either serum-free heavy-RPMI containing 15 µM **PP1** and 10 µM CBR470-1 or serum-free light-RPMI with 10 µM CBR470-1 and DMSO vehicle. Cultures were incubated for a further two hours after which the media was aspirated and both conditions washed with 2 mL fresh 10 µM CBR470-1 containing SILAC RPMI over 40 minutes exchanging the media twice. After washing out excess probe, the cultures were covered with 2 mL SILAC RPMI and irradiated over ice with 365 nm light for 10 minutes. Post-irradiation, samples were processed as described above in the initial PhotoPPI-profile.

**Sample preparation for peroxide perturbed Keap1 PhotoPPI-profile**

Two 10 cm plates seeded with HEK293T cells stably expressing SnapTag fusions of Keap1 were cultured in SILAC RPMI until ~80% confluent. Growth media was removed before administering 150 µM tert-butyl hydrogen peroxide (TBHP) to both heavy and light conditions in 4 mL of full SILAC RPMI and cultures incubated at 37 °C. Six hours later media was removed and replaced with serum-free 150 µM TBHP SILAC RPMI containing either 15 µM **PP1** (heavy) or DMSO (light) then cultures incubated at 37 °C for an additional two hours. Excess probe was removed by aspirating the media then incubating cultures with 2 mL full SILAC RPMI over 40 minutes, exchanging the media twice. Samples were then cover in 2 mL fresh SILAC RPMI and irradiated with 365 nm light over ice for 10 minutes. After irradiation the samples were processed for mass spec analysis as delineated above in the initial PhotoPPI-profile.

**Proteomic LC-MS/MS and data analysis**

LC-MS/MS experiments were performed with an Easy-nLC 1000 ultra high-pressure LC system (ThermoFisher) using a PepMap RSLC C18 column heated to 45˚C (column: 75 µm x 50 cm; 2 µm, 100 Å) coupled to a Q Exactive HF orbitrap and Easy-Spray nanosource (ThermoFisher). Digested peptides (500 ng - 1 µg) in MS/MS Buffer A were injected onto the column and separated using the following gradient of buffer B (0.1% Formic acid acetonitrile) at 300 nL/min: 2-2% buffer B over 5 minutes, 2-25% buffer B over 170 minutes, 25-40% buffer B over 40 minutes, 40-90% buffer B over 10 minutes, 90-90% buffer B over 5 minutes, 90-2% buffer B over 5 minutes, 2-2% buffer B over 5 minutes, 2-90% buffer B over 5 minutes, 90-90% buffer B over 3 minutes, 90-2% buffer B over 5 minutes, 2-2% buffer B over 3 minutes, 2-90% buffer B over 5 minutes, 90-90% buffer B over 5 minutes, 90-2% buffer B over 5 minutes, and 2-2% buffer B over 3 minutes. MS/MS spectra were collected from 0 to 240 minutes using a data-dependent, top 10 ion setting with the following settings: full MS scans were acquired at a resolution of 120,000, scan range of 375-1500 m/z, maximum IT of 60 ms, AGC target of 1e6, and data collection in profile mode. MS2 scans was performed by HCD fragmentation with a resolution of 30,000, AGC target of 1e5, maximum IT of 60 ms, NCE of 27, MSX count of 1, and data type in centroid mode. Isolation window for precursor ions was set to 2.0 m/*z* with isolation offset of 0.0 m/z. Peptides with charge state 1 and undefined were excluded and dynamic exclusion was set to twenty seconds. Furthermore, S-lens RF level was set to 60 with a spray voltage value of 2.20kV and ionization chamber temperature of 275 ˚C.

MS2 files were generated and searched using the ProLuCID algorithm in the Integrated Proteomics Pipeline (IP2) software platform. Human proteome data were searched using a concatenated target/decoy UniProt database (UniProt_Human_reviewed_04-10-2017.fasta). Basic searches were performed with the following search parameters: HCD fragmentation method; monoisotopic precursor ions; high resolution mode (3 isotopic peaks); precursor mass range

600-6,000 and initial fragment tolerance at 600 p.p.m.; enzyme cleavage specificity at C-terminal lysine and arginine residues with 3 missed cleavage sites permitted; static modification of +57.02146 on cysteine (carboxyamidomethylation); two total differential modification sites per peptide, including oxidized methionine (+15.9949); primary scoring type by XCorr and secondary by Zscore; minimum peptide length of six residues with a candidate peptide threshold of 500. A minimum of one peptide per protein and half-tryptic peptide specificity were required. Starting statistics were performed with a Δmass cutoff = 15 p.p.m. with modstat, and trypstat settings. False-discovery rates of peptide (sfp) were set to 1%, peptide modification requirement (-m) was set to 1, and spectra display mode (-t) was set to 1. SILAC searchers were performed as above with "light" and "heavy" database searches of MS1 and MS2 files by including static modification of +8.014168 for lysine and +10.0083 for arginine in a parallel heavy search. SILAC quantification was performed using the QuantCompare algorithm, with a mass tolerance of 10 p.p.m. or less in cases where co-eluting peptide interfere. In general all quantified peptides has mass error within 3 p.p.m..

**Quantitative proteomic data analyses of enriched proteomic samples**

The SILAC ratios of the proteins from enriched proteomic samples were normalized by the median SILAC ratio of the corresponding bulk proteomic sample. The overall normalized SILAC data, from three biologically independent batches and three technical replication LC-MS/MS runs of each batch, were combined. The mean SILAC ratios of each protein were converted to $Log_2$ values, and *P* values were calculated by univariate two-sided t-test with a group of unnormalized SILAC ratios of the protein from enriched samples and a group of median SILAC ratios of the bulk samples. *P* values were further adjusted for Benjamini-Hochberg FDR correction and then converted to $-Log_{10}$ values. The volcano plots were plotted with x-axis of $Log_2$ SILAC values and y-axis of $-Log_{10}$ *P* values (adj.), and the proteins which passed the filter (Probe-to-DMSO ratio > 2, *P* adj. < 0.005) were considered as actual enriched proteins with the PhotoPPI profiling platform.

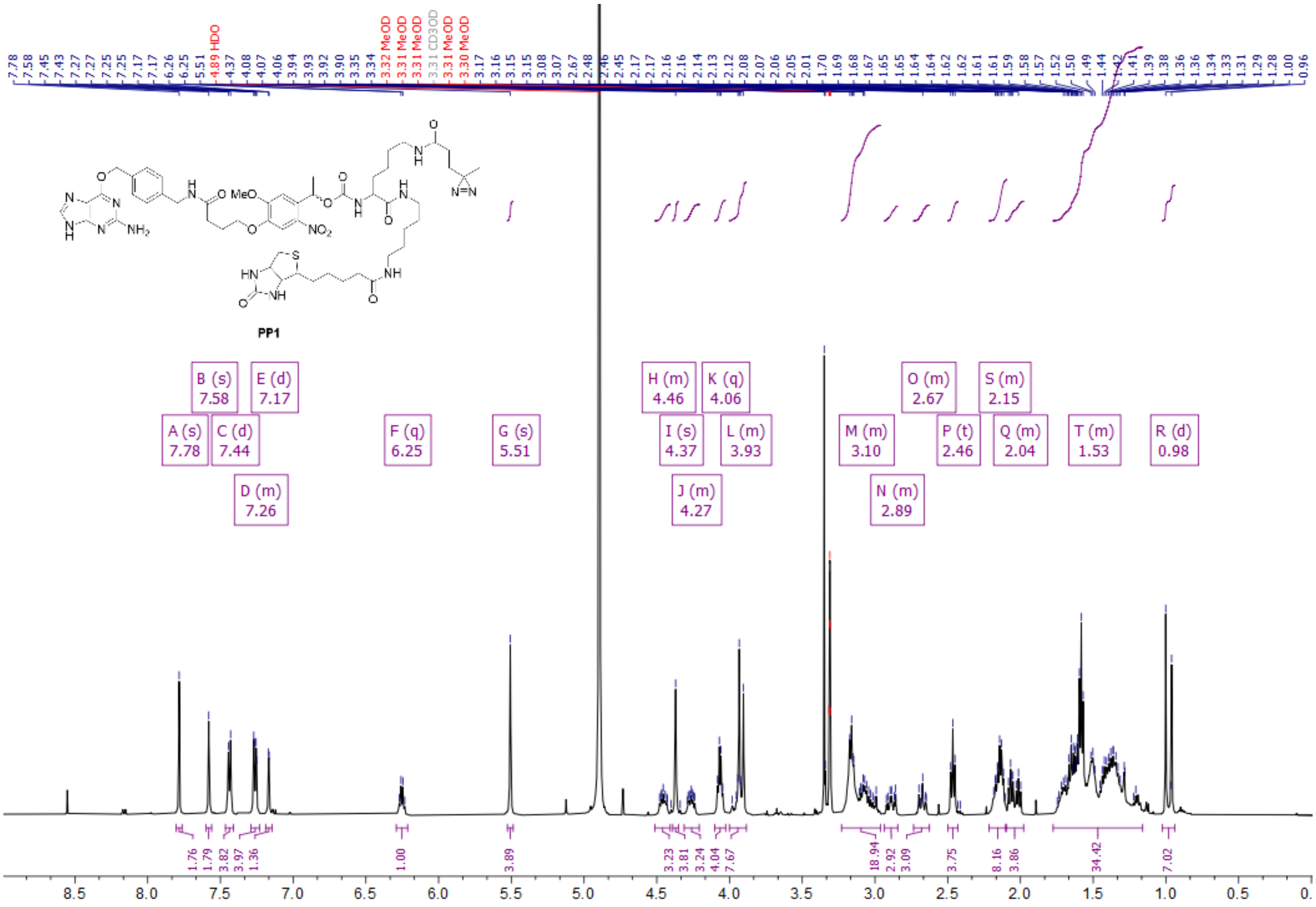
7.78 7.58 7.45 7.43 7.27 7.27 7.25 7.25 7.17 7.17 6.26 6.25 5.51 4.89 HDO 4.37 4.08 4.07 4.06 3.94 3.93 3.92 3.90 3.35 3.34 3.32 MeOD 3.31 MeOD 3.31 MeOD 3.31 CD3OD 3.31 MeOD 3.30 MeOD 3.17 3.16 3.15 3.15 3.08 3.07 2.67 2.48 2.46 2.45 2.17 2.17 2.16 2.16 2.14 2.13 2.12 2.08 2.07 2.06 2.05 2.01 1.70 1.69 1.68 1.67 1.65 1.65 1.64 1.62 1.62 1.61 1.61 1.59 1.58 1.57 1.52 1.50 1.49 1.44 1.42 1.41 1.39 1.38 1.36 1.36 1.34 1.33 1.31 1.29 1.28 1.00 0.96
MeO
NO2
NH2
PP1
A (s) 7.78
B (s) 7.58
C (d) 7.44
D (m) 7.26
E (d) 7.17
F (q) 6.25
G (s) 5.51
H (m) 4.46
I (s) 4.37
J (m) 4.27
K (q) 4.06
L (m) 3.93
M (m) 3.10
N (m) 2.89
O (m) 2.67
P (t) 2.46
Q (m) 2.04
S (m) 2.15
T (m) 1.53
R (d) 0.98
1.76 1.79 3.82 3.97 1.36 1.00 3.89 3.23 3.81 3.24 4.04 7.67 18.94 2.92 3.09 3.75 8.16 3.86 34.42 7.02
8.5 8.0 7.5 7.0 6.5 6.0 5.5 5.0 4.5 4.0 3.5 3.0 2.5 2.0 1.5 1.0 0.5 0.

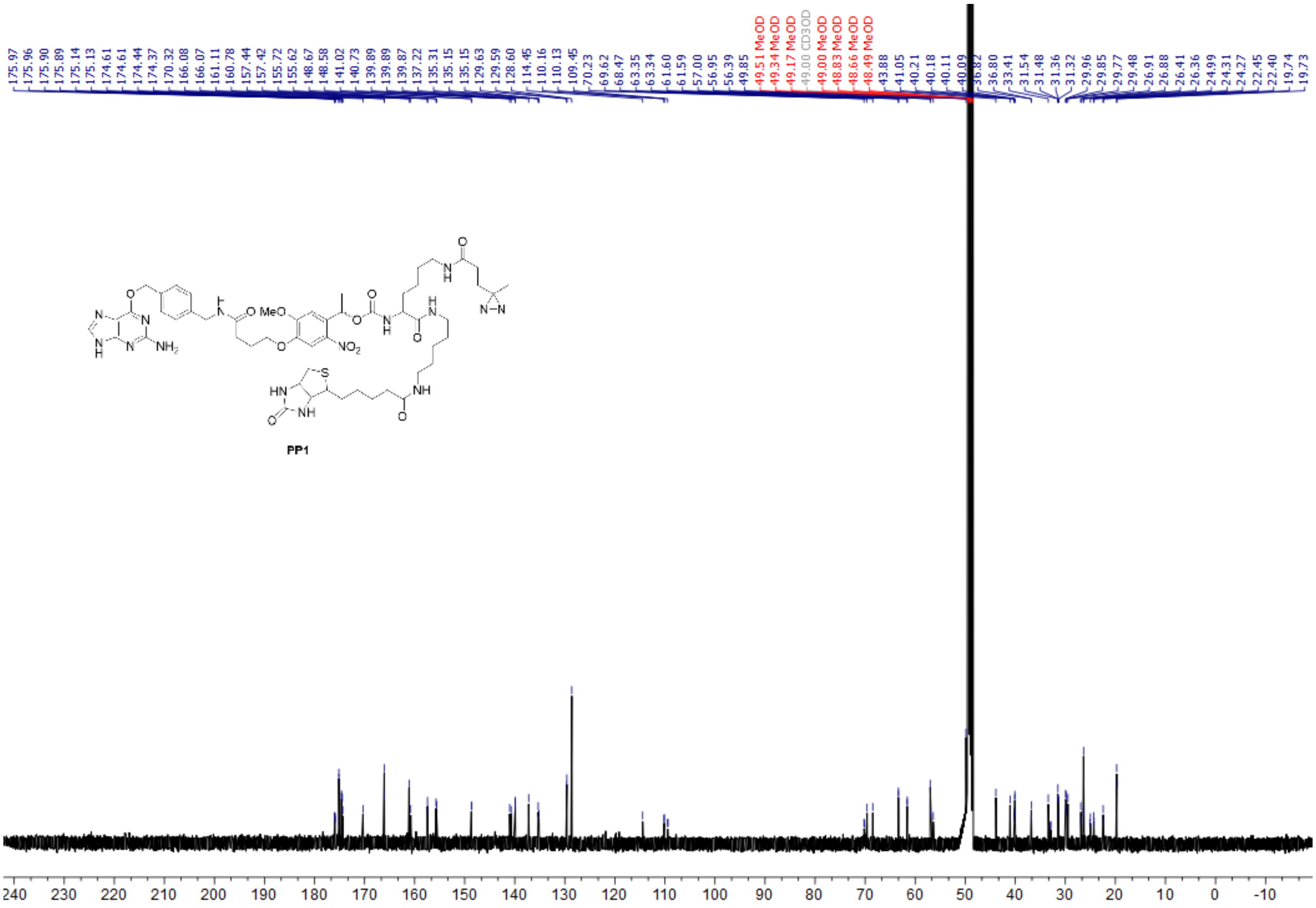
175.97 175.96 175.90 175.89 175.14 175.13 174.61 174.61 174.44 174.37 170.32 166.08 166.07 161.11 160.78 157.44 157.42 155.72 155.62 148.67 148.58 141.02 140.73 139.89 139.89 139.87 137.22 135.31 135.15 129.63 129.59 128.60 114.45 110.16 110.13 109.45 70.23 69.62 68.47 63.35 63.34 61.60 61.59 57.00 56.95 56.39 49.85 49.51 MeOD 49.34 MeOD 49.17 MeOD 49.00 CD3OD 49.00 MeOD 48.83 MeOD 48.66 MeOD 48.49 MeOD 43.88 41.05 40.21 40.18 40.11 40.09 36.82 36.80 33.41 31.54 31.48 31.36 31.32 29.96 29.85 29.77 29.48 26.91 26.88 26.41 26.36 24.99 24.31 24.27 22.45 22.40 19.74 19.73
MeO
NO2
NH2
PP1
240 230 220 210 200 190 180 170 160 150 140 130 120 110 100 90 80 70 60 50 40 30 20 10 0 -10

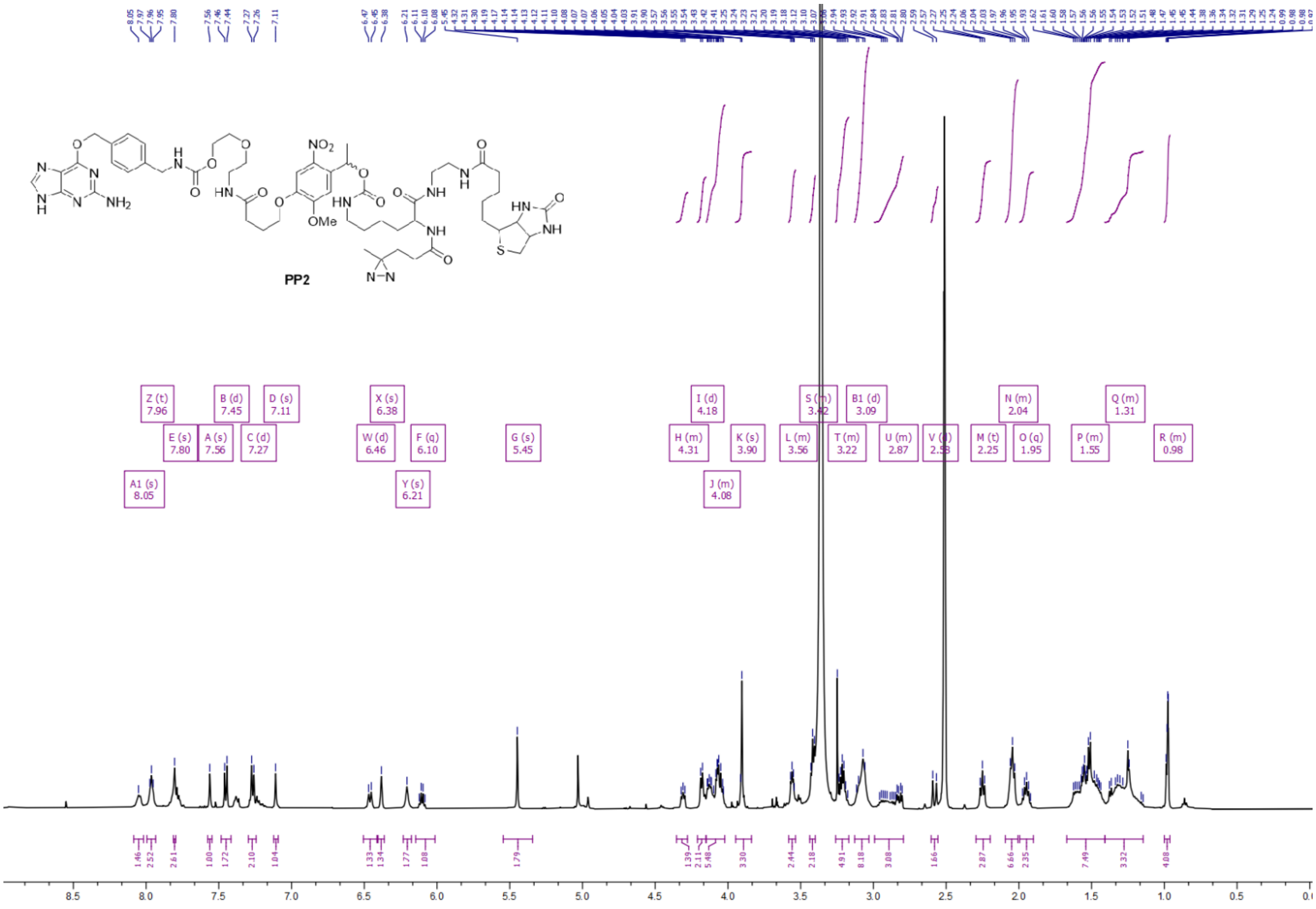
PP2

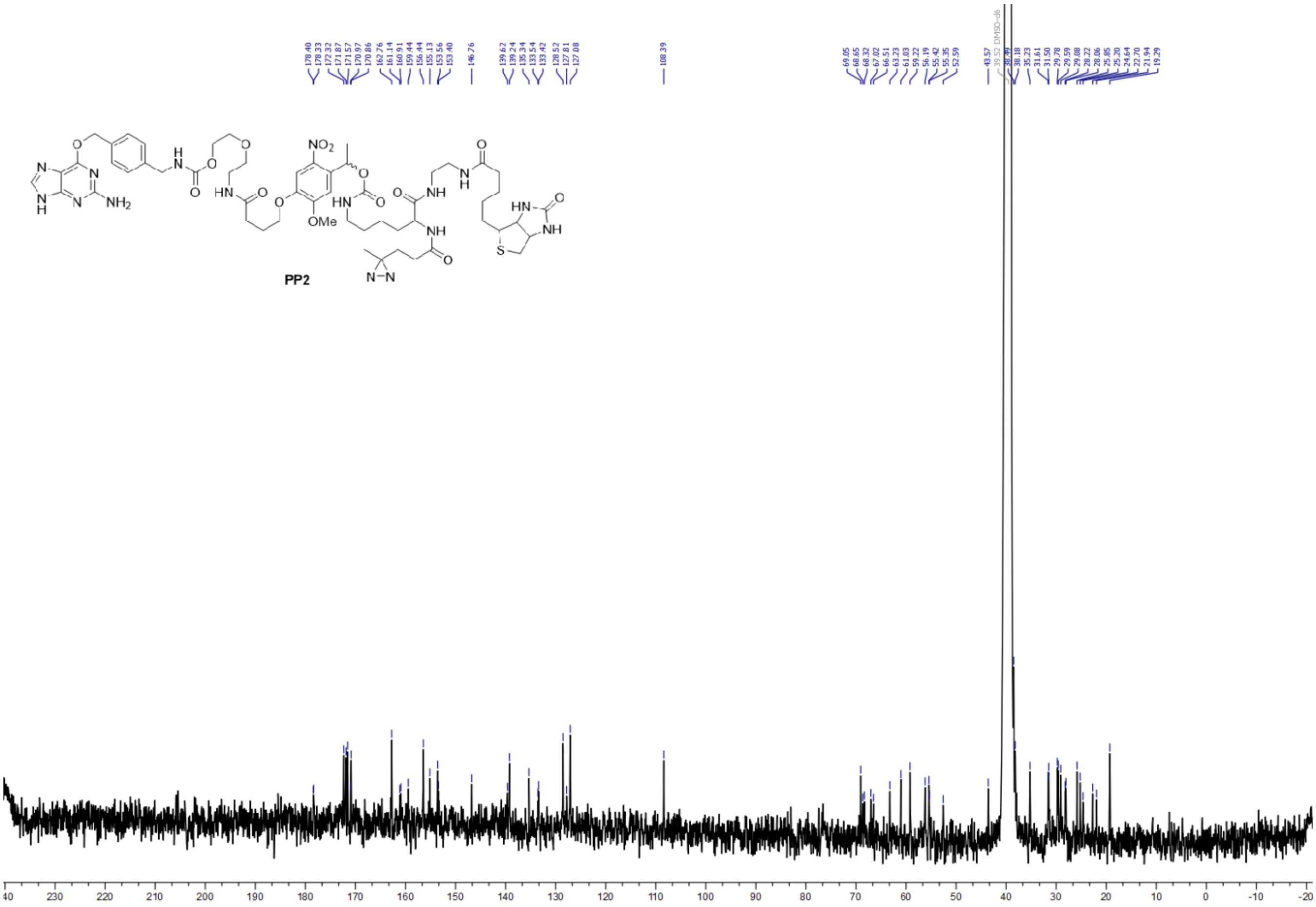
PP2

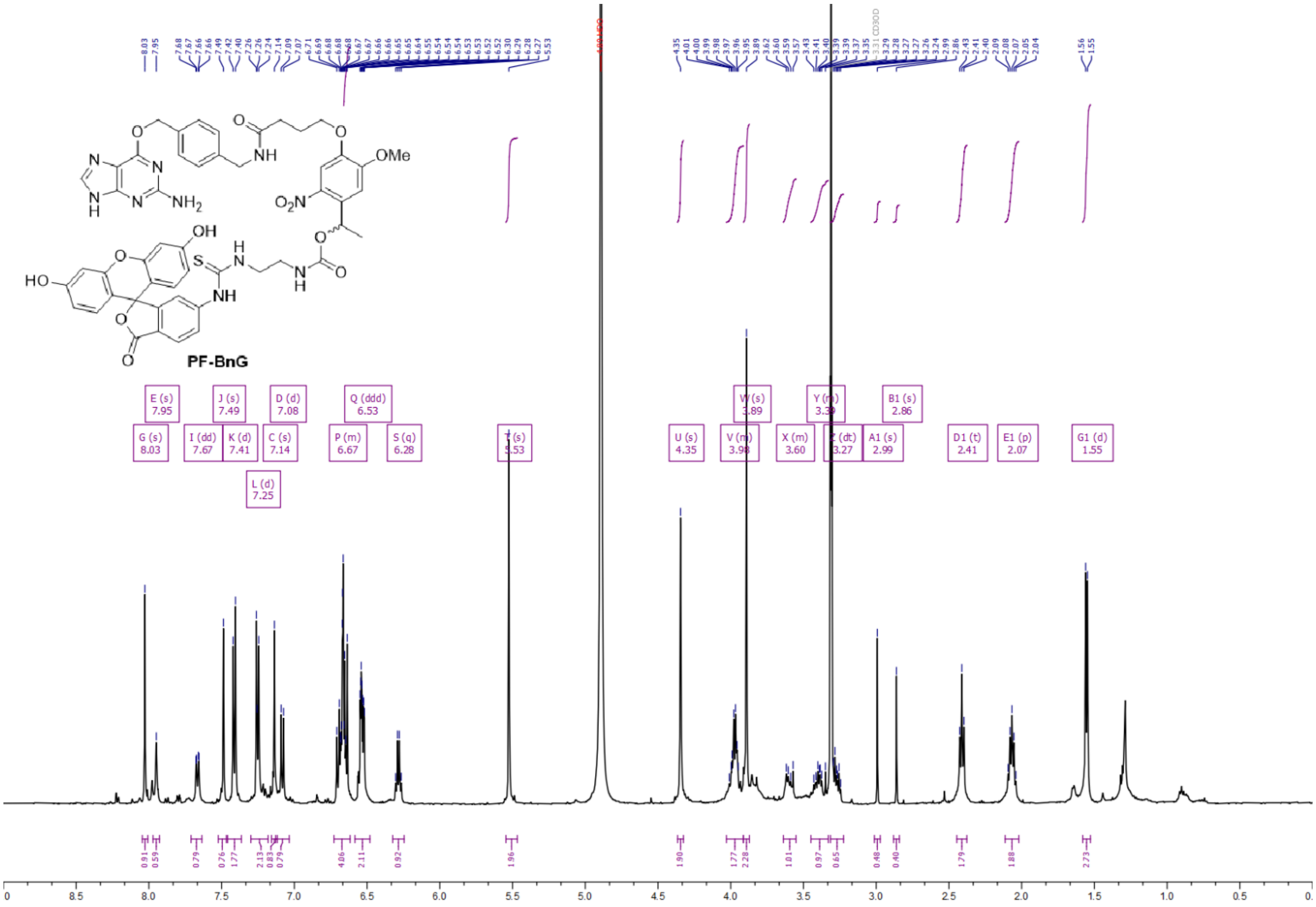
PF-BnG

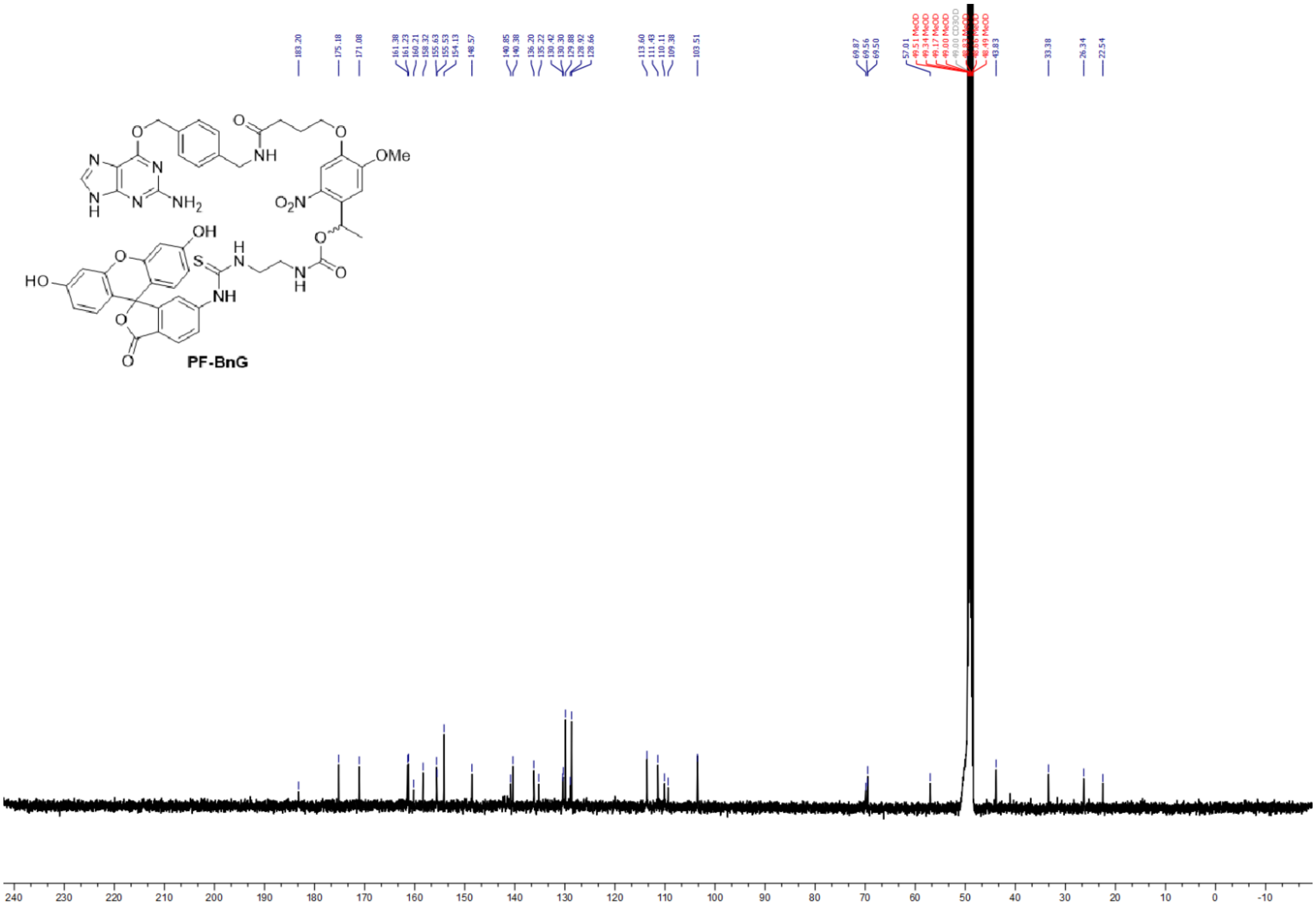
PF-BnG

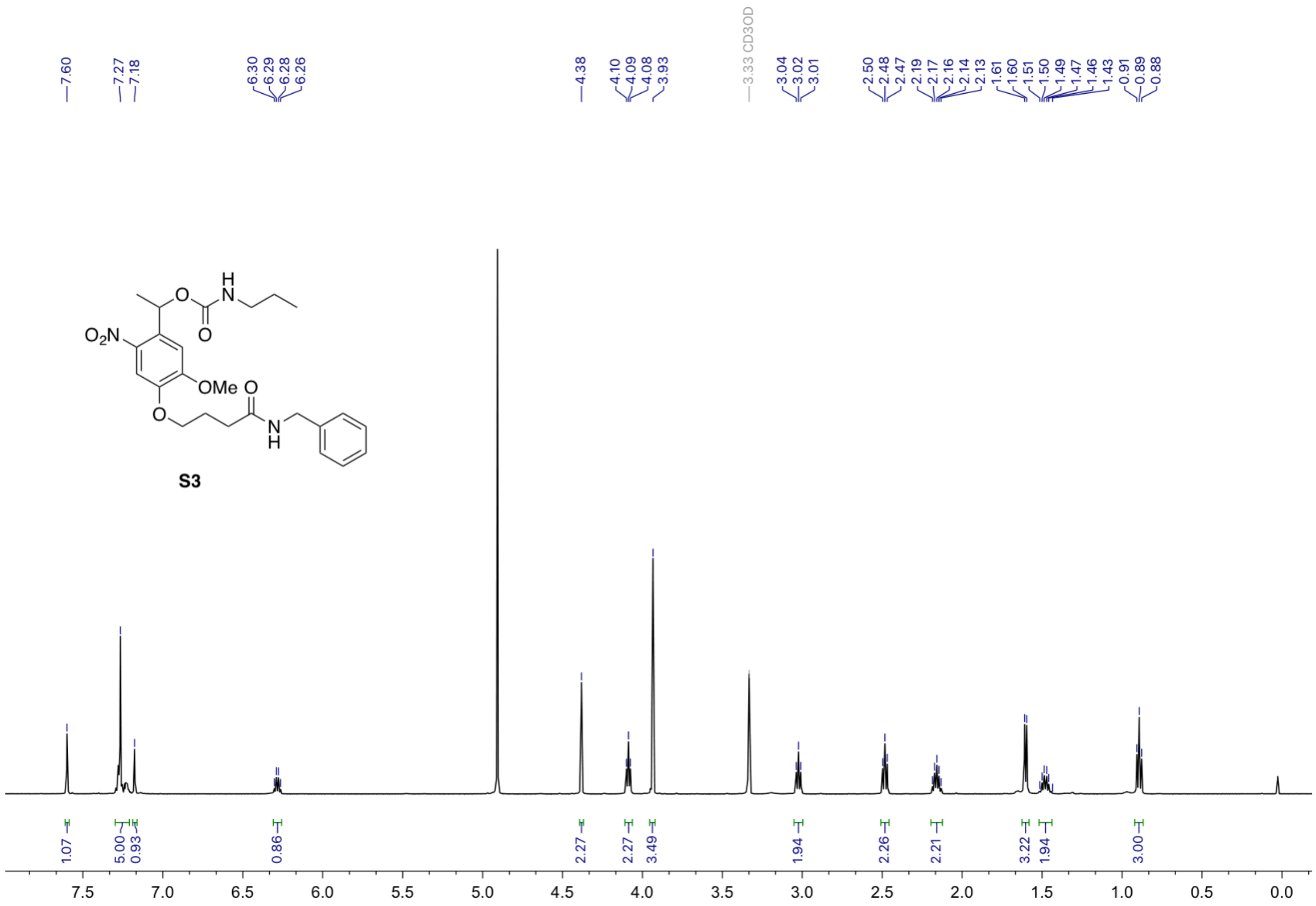

7.60
7.27
7.18
6.30
6.29
6.28
6.26
4.38
4.10
4.09
4.08
3.93
3.33 CD3OD
3.04
3.02
3.01
2.50
2.48
2.47
2.19
2.17
2.16
2.14
2.13
1.61
1.60
1.51
1.50
1.49
1.47
1.46
1.43
0.91
0.89
0.88
O₂N
OMe
S3
1.07
5.00
0.93
0.86
2.27
2.27
3.49
1.94
2.26
2.21
3.22
1.94
3.00
7.5
7.0
6.5
6.0
5.5
5.0
4.5
4.0
3.5
3.0
2.5
2.0
1.5
1.0
0.5
0.0

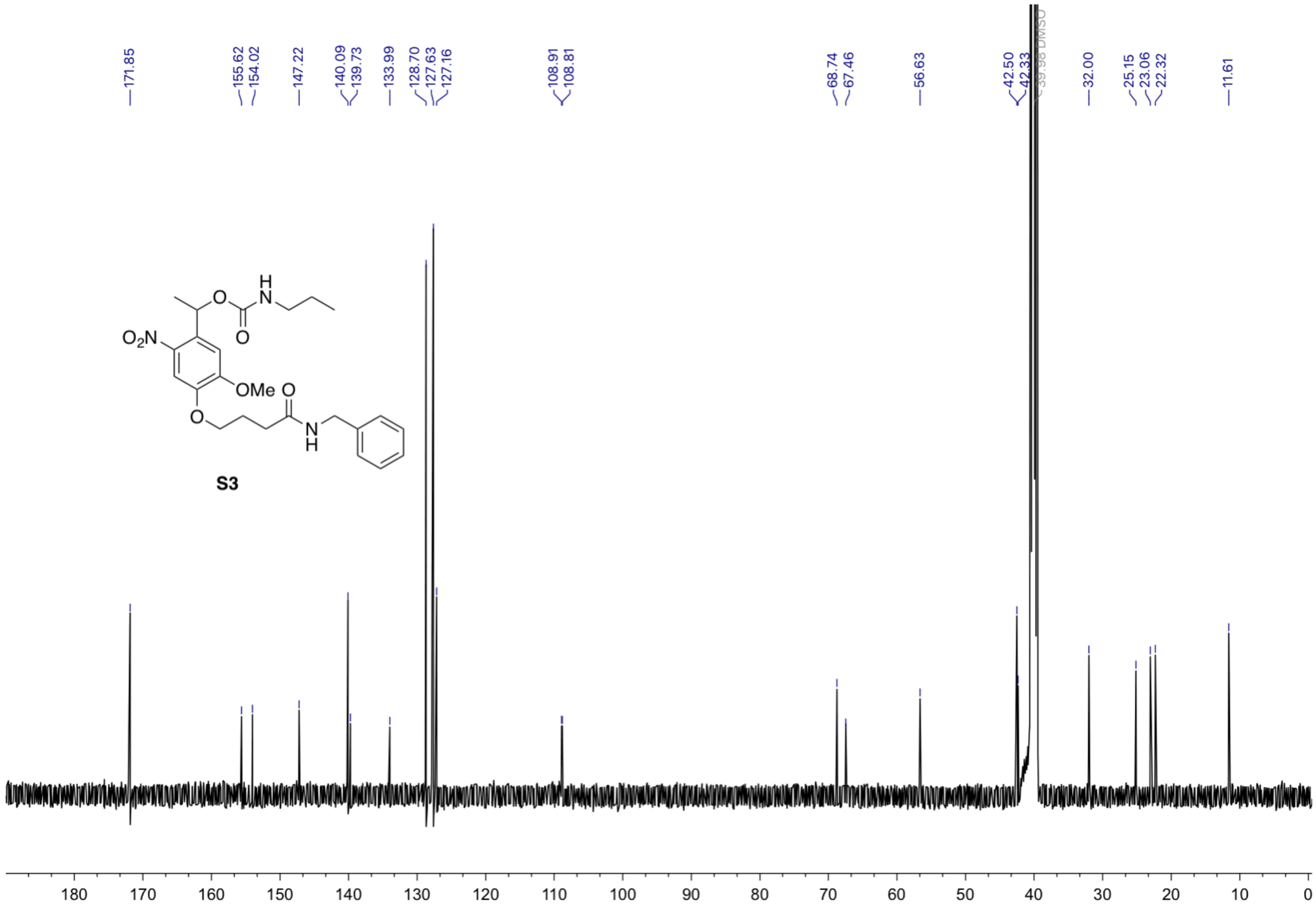
171.85
155.62
154.02
147.22
140.09
139.73
133.99
128.70
127.63
127.16
108.91
108.81
68.74
67.46
56.63
42.50
42.33
39.98 DMSO
32.00
25.15
23.06
22.32
11.61
O
H
N
O
$O_2N$
OMe
O
O
N
H
S3
180
170
160
150
140
130
120
110
100
90
80
70
60
50
40
30
20
10
0
